# DNAJB12 and Hsp70 Facilitate the Conformation Specific Degradation of Arrested N1303K-CFTR Intermediates by ER Associated-Autophagy

**DOI:** 10.1101/2020.10.28.358580

**Authors:** Lihua He, Andrew S. Kennedy, Scott Houck, Andrei Aleksandrov, Nancy L. Quinney, Deborah M. Cholon, Martina Gentzsch, Scott H. Randell, Hong Yu Ren, Douglas M. Cyr

**Affiliations:** Department of Cell Biology and Physiology and the Cystic Fibrosis/Pulmonary Research and Treatment Center, the University of North Carolina at Chapel Hill, Chapel Hill, NC 27599, USA

**Keywords:** Cystic Fibrosis, autophagy, Hsp70, Hsp40, DNAJB12, endoplasmic reticulum, lysosome, proteasome, triage, folding modulator

## Abstract

The transmembrane Hsp40 DNAJB12 and cytosolic Hsp70 cooperate on the ER’s cytoplasmic face to facilitate the triage of nascent polytopic membrane proteins for folding versus degradation. N1303K is the second most common mutation in the ion channel CFTR, but unlike F508del-CFTR, biogenic and functional defects in N1303K-CFTR are resistant to correction bolding modulators. N1303K is reported to arrest CFTR folding at a late stage after partial assembly of its N-terminal domains. N1303K-CFTR intermediates are clients of JB12-Hsp70 complexes, maintained in a detergent soluble-state, and have a relatively long 3-hour half-life. ERAD-resistant pools of N1303K-CFTR are concentrated in ER-tubules that associate with autophagy initiation sites containing WIPI1, FlP200, and LC3. Destabilization of N1303K-CFTR or depletion of JB12 prevents entry of N1303K-CFTR into the membranes of ER-connected phagophores and autolysosomes. Whereas, the stabilization of intermediates with the modulator VX-809 promotes the association of N1303K-CFTR with autophagy initiation machinery. N1303K-CFTR is excluded from the ER-exits site, and its passage from the ER to autolysosomes does not require ER-phagy receptors. DNAJB12 operates in biosynthetically active ER-microdomains to triage in a conformation-specific manner membrane protein intermediates for secretion versus degradation via ERAD or selective-ER associated autophagy.

## INTRODUCTION

Protein triage machines containing Hsp70 sort misfolded proteins between pathways for folding, degradation, and sequestration to suppress the accumulation of toxic species that kill cells through disruption of protein homeostasis (Klaips, Jayaraj et al. 2018). Inherited and spontaneous missense mutations cause misfolding of ion channels, P-Type ATPases, and G protein-coupled receptors, which underlie cystic fibrosis (CF), autoimmune disease, retinitis pigmentosa, hypercholesterolemia, and hypogonadism (Houck and Cyr 2012). CF has been linked to over 1700 different mutations in the CFTR gene, with global misfolding and premature degradation of F508del-CFTR being the most common cause (Veit, Avramescu et al. 2016). Folding modulators help overcome folding defects in F508del-NBD1and, when used in combination with ion channel potentiators, restore CFTR function to therapeutic levels (Donaldson, Pilewski et al. 2018, Keating, Marigowda et al. 2018, Clancy, Cotton et al. 2019). However, it is not clear how changes in the conformation of a membrane protein elicited by a folding modulator would impact its recognition by molecular chaperones and sorting within the ER-membrane system for secretion versus retention and degradation.

Assembly of polytopic membrane proteins is complicated because they expose surfaces that require for folding the coordinated action of cytosolic chaperones such as DNAJA1/Hsp70 (Meacham, Lu et al. 1999), the ER transmembrane chaperones DNAJB12 (JB12) (Li, Jiang et al. 2017), and ER-lumenal BIP and calnexin (Daniels, Kurowski et al. 2003, Houck and Cyr 2012, Behnke, Mann et al. 2016). These same chaperones also facilitate the selection of globally misfolded membrane protein intermediates for ER-associated degradation (ERAD) via the proteasome through interactions with quality control E3 ubiquitin-ligases (Meacham, Patterson et al. 2001) (Cyr, Hohfeld et al. 2002, Younger, Chen et al. 2006, Grove, Fan et al. 2011). How Hsp70 and Hsp40 triage non-native proteins between life and death are not entirely clear, but there are two significant determinations. The first is the folding kinetics for intermediates that are bound and released by Hsp70, with fast folding and limited rebinding to Hsp70 sparing nascent proteins from unnecessary degradation (Qian, McDonough et al. 2006). The second is the cellular set point for the expression of Hsp70 co-chaperones that facilitate folding versus degradation (Meacham, Patterson et al. 2001, Buchberger 2014).

It is generally assumed that globally misfolded proteins are recognized by the Hsp40-Hsp70 system, ubiquitinated by chaperone-dependent E3 ubiquitin ligases, and then threaded into the narrow cavity of the proteasome for degradation (Cyr, Hohfeld et al. 2002, Klaips, Jayaraj et al. 2018). In contrast, aggregated assemblies of misfolded proteins bury surfaces that are recognized by Hsp70 and cannot be translocated into the proteolytic chamber of the proteasomes, so they are degraded by endoproteases localized within autolysosomes (Pohl and Dikic 2019, Wilkinson 2019). However, intermediates of membrane proteins can accumulate in conformations containing tertiary structure {Gautier, 2020 #14760}. Intermediates with stable sub-domains can be difficult to unfold and retrotranslocation into the cytosol and are resistant to ERAD. Still, they are bound by molecular chaperones that suppress their aggregation and protect against ER-stress induced apoptosis (Buchberger 2014, Houck, Ren et al. 2014). ERAD-resistant intermediates of membrane proteins are stabilized by JB12-Hsp70 and degraded by an ER-associated autophagy pathway proposed to involve a functional interplay between JB12-Hsp70 with ER-associated autophagy initiation factors (Houck, Ren et al. 2014). However, the mechanisms that govern the conformation-specific retention and triage of membrane proteins within biosynthetically active regions of the ER-tubular network are not clear.

CFTR is a 1480 amino acid anion channel containing two membrane-spanning domains (MSD1 and MSD2), two nucleotide-binding domains (NBD1 and NBD2), and a disordered regulatory R-domain. CFTR folding and misfolding is the topic of intense study which has identified disease-causing mutations that cause arrested intermediates to accumulate in globally misfolded and aggregated states (Doonan, Guerriero et al. 2019). The F508del mutation is located in NBD1, so F508del-NBD1 is unstable and fails to correctly assemble with NBD2 and CLs transmembrane spans, causing accumulation of globally misfolded F508del-CFTR intermediates. Folding modulators such as VX-809 partially restore F508del-CFTR folding by stabilizing MSD1 and F508del-NBD1, making downstream assembly reactions involving contact formation with CLs more efficient (Ren, Grove et al. 2013, Singh, Fan et al. 2020). In contrast, N1303K, the 2^nd^ most prevalent (2%) CFTR mutation (Cutting 2017), is located in NBD2 and arrests CFTR folding at a late stage, with arrested intermediates of N1303K-CFTR being resistant to repair by folding modulators (Veit, Avramescu et al. 2016). CFTR assembly involves both domain folding and assembly of its NBDs, which occurs through the interaction of NBD1 and NBD2 with each other and with cytosolic loops (CL) located between different pairs of transmembrane helices (TM)(Cyr 2005). NBD2 folding limits completion of CFTR assembly (Zhang, Kartner et al. 1998), so N1303K is likely to disrupt this process. But how molecular chaperones triage and protect the ER from the accumulation of aggregated forms of late-stage intermediates whose folding arrests after initiation of folding is not clear.

To understand how ERQC-machinery triages intermediates membrane proteins a detailed study of the biology and triage of N1303K-CFTR was conducted. These studies lead to the identification of the folding defect in N1303K-CFTR that causes it to stably associate with JB12-Hsp70 and be cleared from the ER membrane system by an ER-associated autophagy mechanism that is not dependent upon ER-phagy receptors (Wilkinson 2019). ER-associated-autophagy is distinct from ER-phagy because ER-tubules containing N1303K-CFTR become clients of autophagy initiation machinery painted on adjacent tubules that convert N1303K-CFTR containing tubules into phagophores containing LC3. Autophagosomes enriched in N1303K-CFTR and LC3 then bud from the ER and fuse with lysosomes that are docked to the WIPI1 rings on which there were formed. These results provide new insights into mechanisms by which Hsp70 and JB12 are associated with ERAD-resistant clients and autophagy initiation machinery to protect the ER from proteotoxic stress.

## RESULTS

### Impact of Folding Modulators and Cl-channel Potentiators on N1303K CFTR Function

To initiate the evaluation of the functional and folding defects caused by N1303K, we measured the response of endogenous N1303K-CFTR expressed in native airway cells obtained from CF patients to folding modulators (Figure 1). Class I folding modulators stabilize MSD1 and NBD1 of CFTR to suppress the folding defects caused by F508del allosterically. Class-II modulators appear to act synergistically with Class I modulators restore assembly of F508del-NBD1 with CL4 in MSD2 of CFTR. (Rosser, Grove et al. 2008, Grove, Rosser et al. 2009, Ren, Grove et al. 2013, Veit, Roldan et al. 2020). Modulators act on arrested intermediates of CFTR but cannot rescue the function of off-pathway and misfolded conformers of CFTR mutants {Van Goor, 2014 #8219}. Therefore, insights into the nature of N1303K-CFTR intermediates are obtained to evaluate their response to modulators.

**Figure 1.**
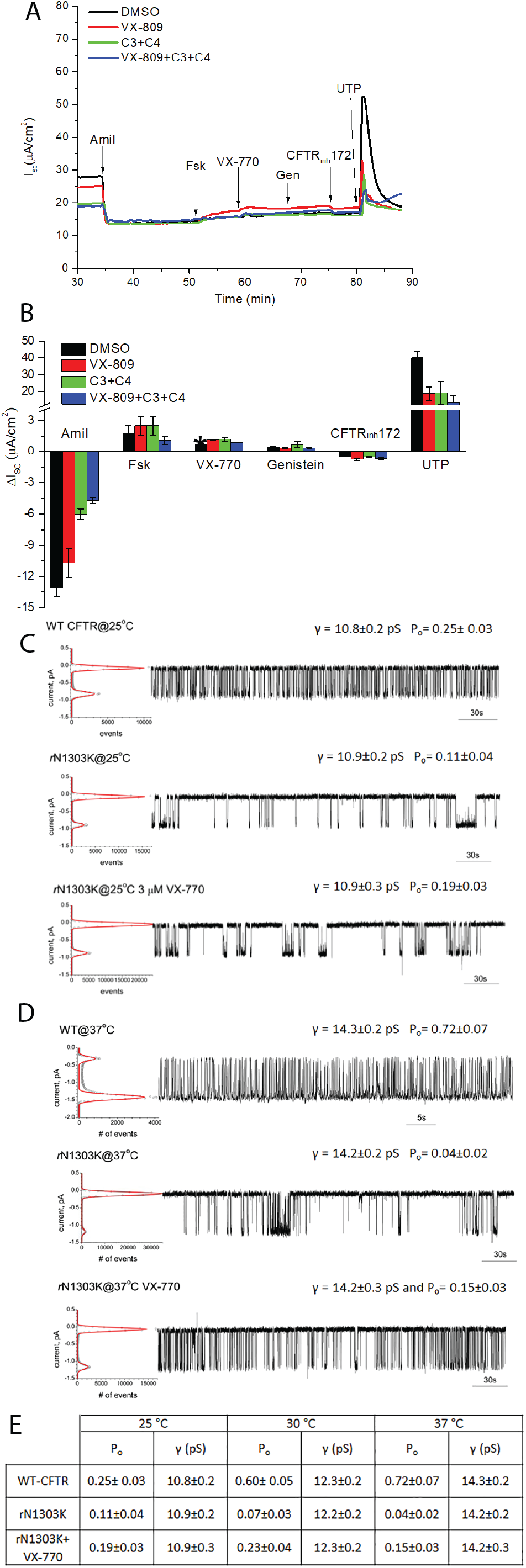
Function of N1303K-CFTR is partially restored by cell culture at low-temperature but is insensitive to folding modulators. (A) The effect of small molecule folding modulators and channel potentiators on CFTR activity in polarized native human bronchial epithelial (HBE) was measured in USSING chambers. Representative traces illustrate the response of CFTR expressed in HBE treated with indicated compounds and CFTR channel activators (B): Bar graphs represent the mean net response (n=4 cultures per corrector treatment) to acutely added stimuli or inhibitors in USSING chamber experiments. HBE from patients carrying N1303K/W1282X mutations were chronically treated for 48 h with 3 μM VX-809, or 5 μM C3+C4, or 0.1% DMSO, as indicated. See Figure S1A for Western blot analysis of the effect of correctors on N1303K/W1282X CFTR expression in HBE. (C) Single channel function of the WT and rescued N1303K (rN1303K) CFTR in the lipid bilayer. The ion channels were transferred into the preformed lipid bilayer by spontaneous fusion of membrane vesicles prepared from BHK cells stably expressing WT or rescued N1303K CFTR. BHK cells expressing N1303K CFTR was rescued by growing at 27 °C in the presence of 5 μM C3 and C4 for the last 24 hrs before harvesting and labeled as rN1303K on the figure. Single channels were recorded at 25 °C (C) or 37 °C (D) for WT CFTR (top panel), rescued N1303K CFTR (middle panel), and rescued N1303K CFTR in the presence of VX-770 (bottom panel). The all points histogram used to calculate single channel conductance (γ) and open probability (Po) is shown on the left of the upper line of each group. The 5 min single channel recording used to prepare all points histogram is shown on the right of the upper line. 7 independent experiments of total 58 minutes duration, 5 independent experiments of total 35 minutes duration, and 4 independent experiments of total 17 minutes duration were used to calculate γ and P_o_ of WT CFTR, rN1303K and rN1303K+VX-770, respectively, at 25 °C (C). 5 independent experiments of total 42 minutes, 4 independent experiments of total 28 minutes duration, and 3 independent experiments with total 12 minutes duration were used to calculate γ and P_o_ of WT CFTR, rN1303K and rN1303K+VX-770, respectively, at 37 °C (D). The difference in conductance between rN1303K and WT CFTR is not significant while the difference in P_o_ is significant (p<0.05). E. Table summarizing the γ and P_o_ of WT and rN1303K CFTR measured at 25, 30 and 37 °C.

In primary airway cells with the heterozygous N1303K/W1282X genotype that was grown on semipermeable-supports at an air-liquid interface to permit the formation of a polarized and sealed cell monolayer, the response of N1303K-CFTR function to modulators was evaluated in USSING chambers (Fulcher, Gabriel et al. 2005). CFTR is a cAMP activated ion channel, so cells were treated with forskolin- to stimulate trans-epithelial short-circuit currents, and N1303K-CFTR function was monitored In the absence or presence of indicated modulators (VX-809, Class I, and C3, C4, Class II) (Gentzsch, Ren et al. 2016, Singh, Fan et al. 2020). W1282X CFTR mRNA is subject to non-sense mediated decay. W1282X CFTR does not accumulate in native airway cells (Oren, Pranke et al. 2017), so the N1303K CFTR channel function is evaluated in the absence of W1282X CFTR in primary N1303K/W1282X cells. N1303K CFTR chloride channel activity in polarized cell monolayers was not detected above the background (Figure 1A, top panel, black trace) and incubation of the cells with VX-809 for 48 hrs. did not significantly increase FSK responses (Figure 1A and B, red trace, and bar graph), nor did a mixture of Class I and Class II modulators (Figure 1A, green trace, and blue trace, respectively). The acute addition of VX-770, a drug that opens the CFTR channel to USSING chambers, caused no significant boost in measured currents. N1303K-CFTR function is, therefore, not detected in the absence of the presence of modulators in primary lung cells. However, modulators appear to impact the conformation of N1303K-CFTR in native airway cells because they increase the steady-state levels of the ER-localized B-form detected by western blot (Figure S1).

Low-temperature cell culture permits some CFTR mutants to fold, escape the ER, and traffic to the plasma membrane, so N1303K-CFTR was stably expressed in BHK cells that were grown at 27 °C. instead of 37 °C {Denning, 1992 #1032}. Single-channel measurements identified N1303K-CFTR channels in detergent-soluble extracts made from the plasma membranes of low-temperature BHK cells (Figure 1C). N1303K-CFTR channels exhibited the same conductance but an open probability (P_o_) of less than half of WT CFTR (Figure 1C, top and middle panels). When the activity of reconstituted N1303K-CFTR channels was measured in single-channel recordings at temperatures from 25 to 37 °C, the open probability (P_o_) of WT CFTR increased to 0.72. It remained stable during the time course of measurements (Figure 1D, top trace). However, the P_o_ for N1303K CFTR decreased (Figure 1D, middle trace), and channel activity was less stable than WT-CFTR at 37 °C. A summary of the P_o_ and conductance measured at different temperatures is presented in Figure 1E. Notably, the P_o_ of N1303K CFTR was stabilized by VX-770 at 25, 30, and 37 °C. These results suggest that N1303K-CFTR can assemble into a CFTR-like ion channel when thermal energy is low. These data indicate that N1303K-CFTR intermediates’ fate can be altered to suppress recognition of them by ERQC factors.

### N1303K-CFTR intermediates accumulate in a detergent-soluble and biochemically stable state

To determine the challenge presented to the ERQC-machinery by late-stage intermediates of membrane proteins that fail to complete folding, we characterized the behavior of N1303K-CFTR in ER membranes of cells cultured at 37 °C (Meacham, Lu et al. 1999, Meacham, Patterson et al. 2001, Younger, Chen et al. 2006, Ren, Grove et al. 2013, Gentzsch, Ren et al. 2016). N1303K-CFTR intermediates accumulated in an incompletely glycosylated state in the ER that is termed band B, that has faster electrophoretic mobility than the folded form, represented by band C. Band C is glycosylated further during passage through the Golgi on route to the plasma membrane (Figure 2) (Meacham, Lu et al. 1999, Meacham, Patterson et al. 2001, Dalal, Rosser et al. 2004, Cyr 2005). Levels of the B-form of N1303K-CFTR were several-fold higher than the B-form of F508del-CFTR (Figure 2A and Figure S2). These differences were accounted for by the observed longer half-life of N1303K-CFTR (3-hr) versus F508del-CFTR (1-hr) in cycloheximide (CHX) chase experiments (Figure 2A). A short 2-hr pre-treatment with VX-809 (3 μM) further increased the stability of N1303K-CFTR such that at the end of 3-hr CHX-chase, around 80% of B-band remained. VX-809 treatment increased the accumulation of the folded C-form of ΔF508-CFTR but only increased the half-life of its B-form to 1.5 hr. N1303K-CFTR accumulates in a conformation that is stabilized by VX-809. Yet, at physiological temperatures, this Class I modulator does not promote the assembly of N1303K-CFTR into channels that escape the ER.

**Figure 2.**
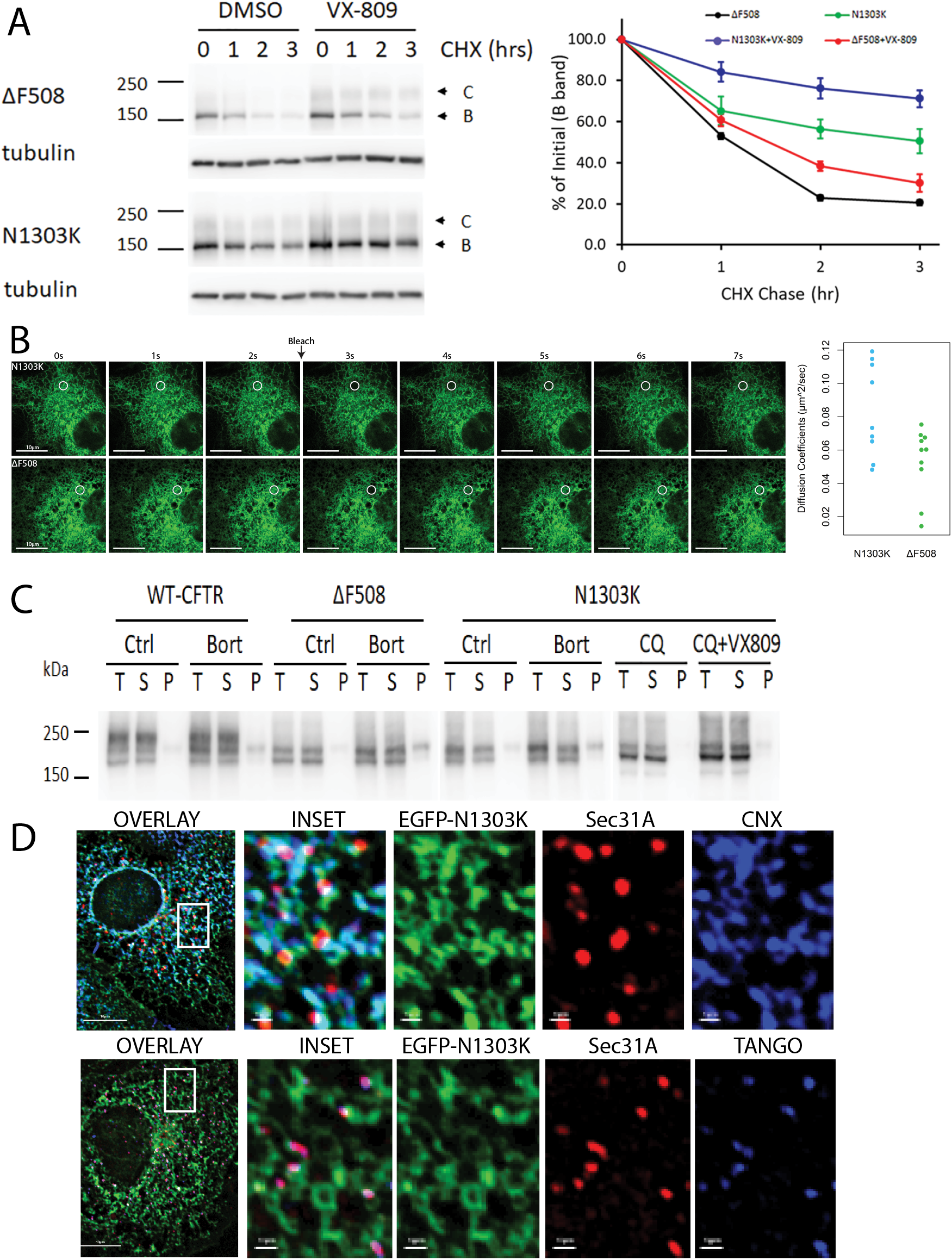
N1303K CFTR is more stable than ΔF508 CFTR and is partially degraded by autophagy. (A). Cycloheximide (CHX) chase of N1303K and ΔF508 CFTR in the absence or presence of VX-809. HEK cells were transiently transfected with N1303K and ΔF508 CFTR. At 24 hrs. after transfection, cells were pretreated with or without 5 μM VX-809 for 2 hrs., followed by addition of 100 μM CHX and incubation for the indicated time. Cell lysates in SDS-PAGE sample buffer were subjected to 7.5% SDS-PAGE and Western blot analysis with CFTR mAb 596. Protein loading was normalized by reprobing the blots with anti-tubulin antibody. Representative blots are shown on the left, with quantification normalized to tubulin shown on the right (n=3). CHX chase of N1303K and ΔF508 CFTR after overnight VX-809 treatment gave similar results. (B). Live-Cell imaging and FRAP of EGFP-N1303K CFTR and EGFP-F508del CFTR that was transiently expressed in COS-7 cells and imaged with a Zeiss-880 microscope using a PMT detector. The swarm plot shows the diffusion coefficients for N1303K (8 trials) and ΔF508 CFTR (10 trials). (C). Fractionation of WT, F508del and N1303K CFTR into Triton X-100 soluble and insoluble fractions of cell extracts. (D) N1303K is excluded from ER exit sites and co-localizes in the tubular ER with calnexin. COS-7 cells were transiently transfected with N1303K CFTR. 18 hrs. after-transection and 6 hrs. prior fixation cells were treated with 15 μM chloroquine (CQ). Immunostaining was carried out on methanol fixed cells with anti-GFP, and with antibodies to endogenous Sec31A and calnexin (CNX)

To study the intracellular behavior of N1303K-CFTR via microscopy, GFP-N1303K-CFTR and GFP-F508del-CFTR were employed as tools in imaging (Figure 2–7). In controls, untagged and GFP-tagged CFTR behaved similarly (Figure 3A versus 2, 3, 4-7)), so triage of GFP-N1303K-CFTR and GFP-F508del reflects the behavior of untagged CFTR. In the characterization of CFTR intermediates, the following was observed (Figure 2B-C): 1, F508del-CFTR, and N1303K-CFTR accumulate in apparently healthy tubular regions of ER in the absence of visible aggregation. 2. In live cells, GFP-F508del-CFTR and GFP-N1303K-CFTR accumulate within dynamic ER tubules, and both diffuse at similar rates into photobleached regions. 3. The ER-localized B-form of F508del-CFTR and N1303K-CFTR do not form detergent-insoluble aggregates that partition into the pellets of centrifuged cell extracts prepared with the detergent Triton-X-100 (Figure 2C). 4. The modulator VX-809, while increasing the half-life of N1303K CFTR, did not lead it to accumulate in the detergent-insoluble fraction of cells. Also, inhibition of the proteasome (bortezomib (Bort)) or lysosomal proteases (chloroquine (CQ)) for 6-hours, which enhances the accumulation of the B-form, does not lead N1303K-CFTR to enter into detergent-insoluble aggregates. 5. N1303K-CFTR accumulates in ER tubules containing the molecular chaperone calnexin but is excluded from ER exit sites that are marked by TANGO1 and the vesicle coat protein Sec31 (Ge, Zhang et al. 2017) (Figure 2D). Overall, N1303K-CFTR is reported to accumulate in the ER in a conformation that can be stabilized by VX-809, but N1303K intermediates cannot fold, and these kinetically trapped intermediates persist within ER membranes.

**Figure 3.**
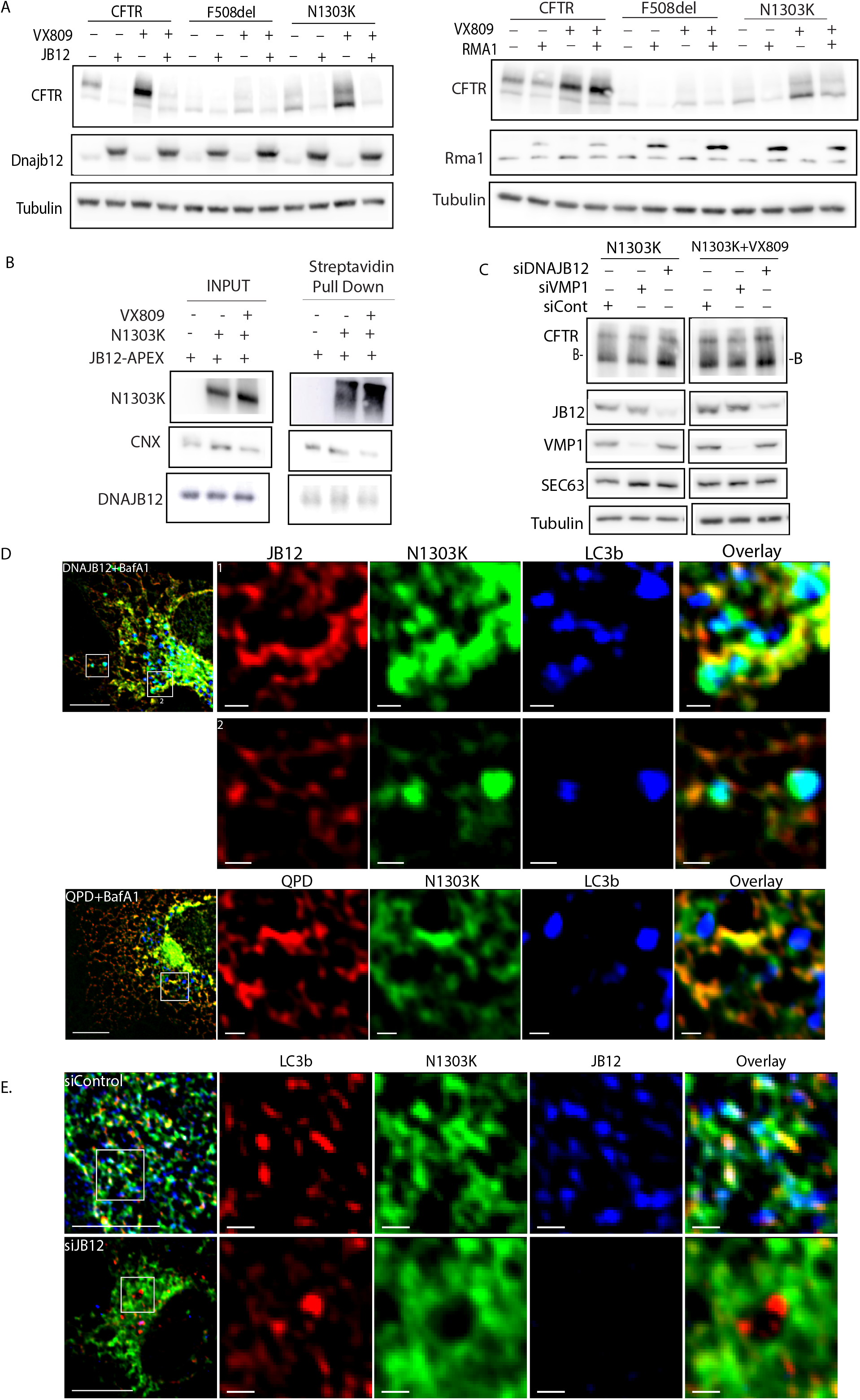
N130K-CFTR is sensitive to elevation of JB12 and RMA1/RNF5 and accumulates in ER tubules with JB12 or membranes that exclude JB12 and contain LC3B. (A) Changes in steady-state levels of CFTR, F508del-CFTR, and N1303K-CFTR in the absence or presence of VX-809 that are caused by the elevation of JB12 or RMA1/RNF5. Indicated forms of CFTR were expressed in the absence or presence of VX-809 that was added to cell cultures 6-hours prior to harvest and analysis by western blot. Western blots in middle panels show levels of overexpressed JB12 (50 ng of pCDNA3-DNAJB12) or RMA1/RNF5 (100 ng of pCDNA-RMA1/RNF5) relative to endogenous pools. (B) Proximity labeling of N1303K-CFTR by JB12-APEX in the absence or presence of VX-809. JB12-APEX, which contains an APEX domain localized in the ER-lumen was co-expressed with N1303K-CFTR in the absence or presence of VX-809. Cells were incubated with hydrogen peroxide and biotin-phenol and modified adducts were then isolated from cell extracts by streptavidin-pull downs and detected by western-blot with indicated antibodies. (C) Analysis of the localization of GFP-N1303K-CFTR in the ER relative to JB12 or RMA1/RNF5. N1303K-CFTR was expressed in COS-7 cells in the absence or presence of JB12 (expressed 50 ng of pCDNA3). Cells were fixed with methanol and stained for JB12 or endogenous LC3. Immunofluorescent images were collected with an Olympus IX-81 with CellSens software and processed with ICY. (D) Overexpressed FLAG-DNAJB12 or FLAG-QPD-DNAJB12 with GFP-N1303K-CFTR in COS-7 cells. Cells were fixed in methanol at −20°C for 5 minutes, then blocked for 1 hour at ambient temperature with blocking buffer (3% bovine serum albumin in phosphate buffered solution), followed by primary and secondary antibody labeling diluted with blocking buffer. After washes with blocking buffer, coverslips were mounted using ProLong Diamond with DAPI mounting solution, and sealed immediately with nail polish. (E) siRNA (or control) for DNAJB12 and transiently transfected (effectene) with GFP-N1303K in COS-7 cells, then fixed (using same procedure as in D).

### N1303K-CFTR Segregates into ER Sub-domains with the Hsp40 DNAJB12 or the Autophagy Factor LC3B

We explored why N1303K-CFTR has a 3-hr half-life versus 1-hr. for F508del-CFTR by examining the respective intermediates’ sensitivity to modulation of the ERAD factors JB12 and RMA1/RNF5 (Figure 3A). Accumulation of N1303K-CFTR and F508del-CFTR was sensitive to the elevation of JB12 or RMA1, and the addition of VX-809 did not protect significant portions of intermediates from JB12 or RMA1. In proximity studies, APEX-JB12, which has an ER luminal APEX, labeled N1303K-CFTR in the absence and presence of VX-809 (Figure 3B). Also, the siRNA depletion of JB12 was accompanied by an increase in the steady-state levels of N1303K-CFTR. In contrast, a control depletion of an unrelated transmembrane VMP1, which regulates the SERCA2B function, did not impact steady-levels of N1303K-CFTR (Figure 3C). N1303K-CFTR is similar to WT-CFTR and F508del-CFTR in that it is a client of JB12 (Grove, Fan et al. 2011). Even though VX-809 stabilizes N1303K-CFTR, VX-809 treatments do not reduce proximity labeling of N1303K-CFTR by JB12-APEX or decrease the sensitivity of N1303K-CFTR to modulation of JB12 or RMA1 activity. N1303K-CFTR intermediates appear to become kinetically trapped, and even though VX-809 increases the accumulation of trapped intermediates, they remain clients of JB12-containing complexes.

Misfolded intermediates of membrane proteins that have long half-lives can be resistant to ERAD. Those that do not form large detergent-insoluble aggregates are cleared from the ER via a selective ER-associated autophagy mechanism (Houck, Ren et al. 2014). To evaluate the triage of N1303K-CFTR, we explored via fluorescence microscopy the localization of its intermediates in the ER membrane system with JB12 and the autophagy initiation factor LC3B (Figure 3D). Remarkably, GFP-N1303K-CFTR was found in two different subsets of interconnected ER-tubules containing either overexpressed WT-JB12 or endogenous LC3B (Figure 3D). The inset shows JB12 (red) localized in tubules with GFP-N1303K-CFTR (green) that form a perimeter around an adjacent region of the ER enriched in GFP-N1303K-CFTR and LC3B. LC3 is conjugated to phosphatidylethanolamine within the membranes of ER-connected phagophores at initial phases of autophagy (Yla-Anttila, Vihinen et al. 2009, Matsunaga, Morita et al. 2010). Membranes containing N1303K-CFTR may segregate within the ER into microdomains known as omegasomes where ER-tubules are modified and converted into phagophores {Matsunaga, 2010 #6196;Matsunaga, 2010 #6196;Yla-Anttila, 2009 #11134;Axe, 2008 #4587;Ktistakis, 2020 #6477;Zachari, 2019 #5144}. Also, we detected co-localized signals for GFP-N1303K-CFTR and LC3B in non-ER attached foci that resemble autolysosomes.

QPD-JB12, containing a mutation in the J-domain’s HPD motif, binds clients but cannot release them because QPD-JB12 can no longer interact with Hsp70 (Grove, Fan et al. 2011, Sopha, Ren et al. 2017). QPD-JB12 co-localized with GFP-N1303K CFTR in the ER and hindered the segregation of N1303K-CFTR into ER-membranes decorated with LC3B. Hsp70 is not required for basal autophagy, and QPD-JB12 expression does not interfere with the formation of autophagic foci containing LC3B (Figure 3D). QPD-JB12’s inability to interact with Hsp70 and cycle on and off N1303K-CFTR, therefore, appears to selectively inhibit the segregation of N1303K-CFTR into ER-membranes decorated with LC3B.

Endogenous JB12 was also detected via microscopy in ER microdomains with GFP-N1303K-CFTR and endogenous LC3B (Figure 3E). ER-tubules containing endogenous JB12 and GFP-N1303K-CFTR intersect with tubules decorated with endogenous LC3B that extend from out from ER-rings. The depletion of JB12 by siRNA was accompanied by an expansion of N1303k-CFTR containing ER-membranes and the exclusion of N1303K-CFTR from LC3B-decorated ER tubular extensions.

In summary, kinetically-trapped intermediates of N1303K-CFTR are present in complexes containing JB12. The depletion of JB12 causes the expansion of ER-domains containing N1303K-CFTR and hinders N1303K-CFTR segregation into LC3B-decorated ER-tubules. JB12 depletion/inactivation does not reduce the formation of autophagic foci containing LC3B, so it appears to be required for the selective autophagic degradation of N1303K-CFTR.

### N1303K-CFTR intermediates, but not F508del-CFTR, traffic from the ER to Autolysosomes

How do the N1303K-CFTR co-localize with ER-tubular extensions and decorated with LC3B transit to autolysosomes (Figure 4)? To address this question, we first, we repeated experiments conducted with GFP-N1303K-CFTR with untagged CFTR and demonstrated that N1303K-CFTR, but not F508del-CFTR, accumulates in autophagic foci whose central core contained LC3B (Figure 4A). Three-color imaging showed that GFP-N1303K-CFTR and LC3B positive puncta also have LAMP1/2, so they are, in fact, autolysosomes (Figure 4B).

**Figure 4.**
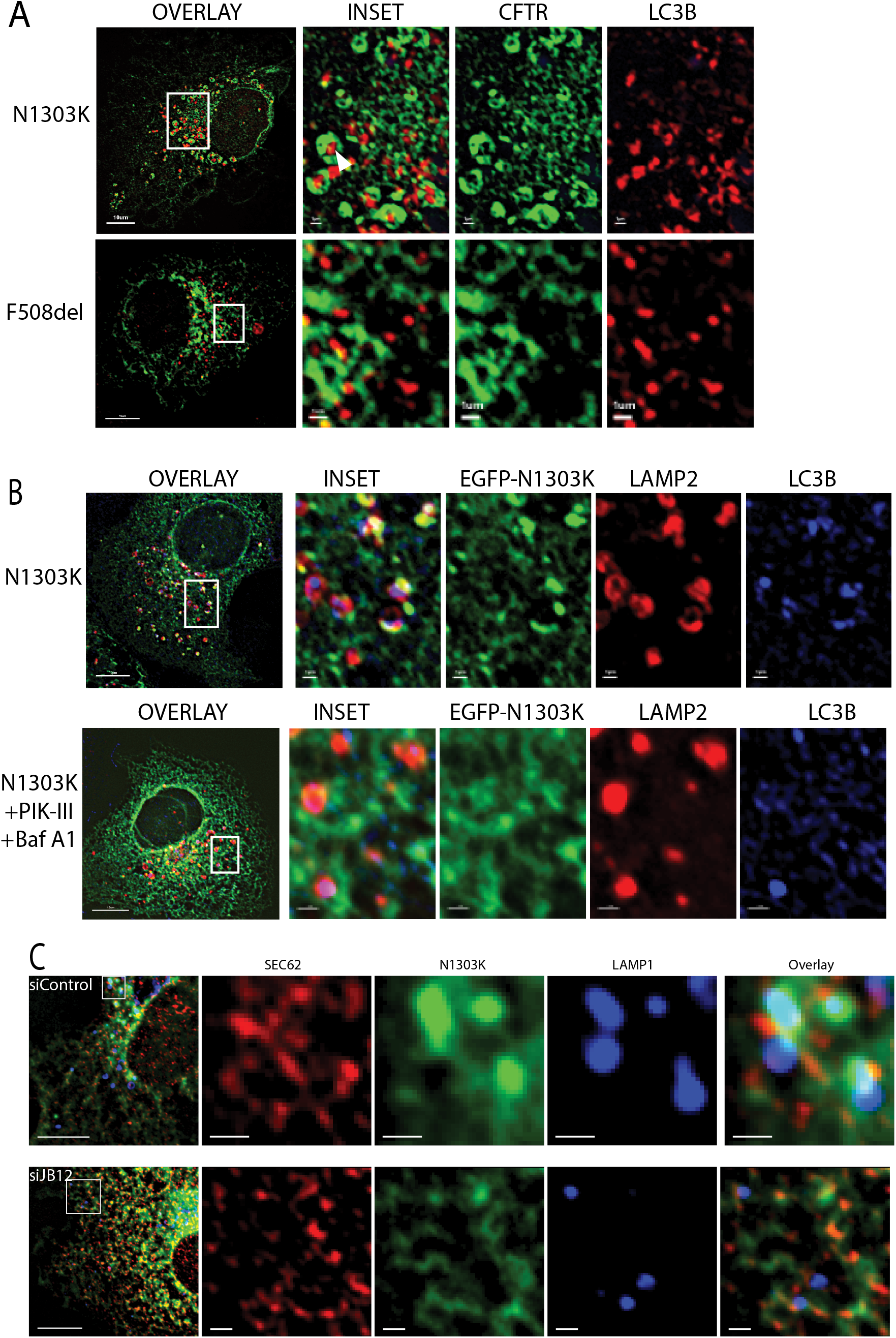
JB12 is Necessary for Traffic of N1303K-CFTR from the ER to Autolysosome. (A) Localization in COS-7 cells of untagged N1303K-CFTR, but not F508del-CFTR, in foci with endogenous LC3B. (B) Localization of GFP-N1303K-CFTR in autolysosomes with endogenous LAMP1 and LC3B is sensitive to PI3K-III, an inhibitor of the autophagy initiation kinase, VPS34. (C) Depletion of JB12 from COS-7 cells by siRNA reduces movement of N1303K-CFTR from the ER to LAMP1 positive foci. JB12 levels were reduced via siRNA and 18-hours later cells were transfected with GFP-N1303K-CFTR. Cells were fixed 18 hours after introduction of GFP-N1303K-CFTR and stained for endogenous LAMP1 or the ER-marker Sec62. Immunofluorescent images were collected with an Olympus IX-81 with CellSens software and processed with ICY.

The autophagy initiation kinase (VPS34) inhibitor PIK-III blocked the entry of GFP-N1303K-CFTR into autolysosomes containing LAMP1 (Figure 4B) and did so under conditions where conversion of LC3B-I to LC3B-II was abolished (Figure S3B). LC3B II being the form that is covalently attached to the phosphatidylethanolamine headgroup of ER-lipids during initiation of phagophore formation. Likewise, siRNA of Beclin-1, treatment of cells with a ULK1 inhibitor, and overexpression of a dominant-negative form of ULK1 also blocked the transit of GFP-N1303K-CFTR from the ER to autolysosomes (Table S1). Importantly, siRNA depletion of JB12 caused GFP-N1303K-CFTR to accumulate within biosynthetically active regions of the ER containing protein translocon associated protein Sec62 reduced co-localization LAMP1 (Figure 4C). Sec62 and JB12 each co-localize in the ER with N1303K-CFTR, but Sec62 and JB12 are not detected with GFP-N1303K-CFTR in membranes decorated with LC3B or in autolysosomes with LAMP1. N1303K-CFTR intermediates appear separate from JB12 in a process requiring Hsp70 before entry into ER membranes containing LC3B and the subsequent passage from the ER to autolysosomes.

### ER Membranes Enriched in N1303K-CFTR Protrude from WIPI1 Rings Containing Docked Autolysosomes

The route taken by N1303K-CFTR from the tubular-ER to the autolysosome was explored by visualizing its association with the omegasome factors DFCP1 and WIPI1, and LAMP1. The VPS34 kinase complex produces pools of inositol-3-phosphate head groups on membrane lipids bound by cytosolic DFCP1 and WIPI1, which facilitate downstream reactions phagophore formation involving LC3 conjugation and autophagosome biogenesis {Tooze, 2010 #4416}. GFP-N1303K-CFTR co-localizes in tubular ER-membranes with endogenous pools of the Sec62 and mCherry-DFCP1 (Figure 5A). Similarly, GFP-N1303K-CFTR is detected with mCherry-WIPI1 in domains containing the ULK1 subunit FIP200 (Figure 5B). Interestingly, GFP-N1303K-CFTR containing ER-tubules extend out from the surface of WIPI1 and FIP200 decorated ER membranes in apparent phagophores. Before exiting from the ER, it appears that N1303K-CFTR is concentrated in ER-tubules that associate with ER membranes that are decorated with FIP200, which controls ULK1 localization and thereby defines sites for focal activation of selective autophagy (Vargas, Wang et al. 2019). Use of STED microscopy to image GFP-N1303K-CFTR, LC3-mCherry, and BFP-LAMP1 in live cells enabled visualization of GFP-N1303K and LC3 containing foci that interact with autolysosomes containing BFP-LAMP1 (Figure 5C).

**Figure 5.**
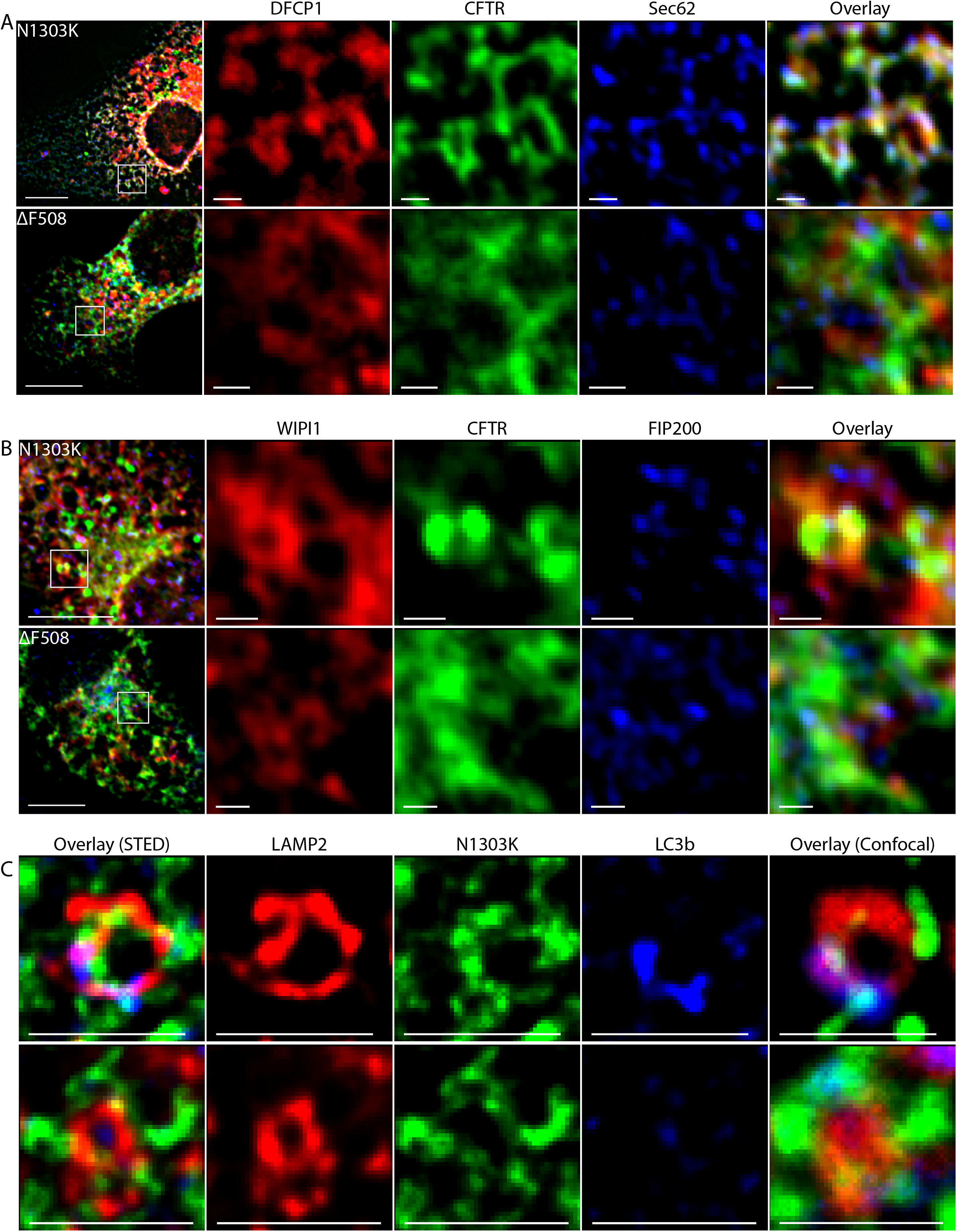
N1303K-CFTR intermediates, but not F508del-CFTR, are detected by fluorescence microscopy with autophagy initiation machinery during sorting within the ER and trafficking to autolysosome. (A) N1303K-CFTR co-localizes in ER-tubules with the autophagy initiation factor mCherry-WIPI1 and the translocon associated Sec62. (B) ER-attached membranes containing N1303K-CFTR extend from rings of ER-membrane that is decorated with mCherry-WIPI1 and endogenous FIP200. (C) Super-resolution analysis of associations between N1303K-CFTR, RPF-WIPI1 and BFP-LC3 in live COS-7 cells. A and B display images of fixed COS-7 cells expressing GFP-N1303K-CFTR, and the indicated mCherry-tagged or endogenous marker proteins obtained with an Olympus IX-81 and CellSens software. In images displayed in C were obtained from fixed cells at super-resolution with a Lecia STED Microscope.

Data presented are interpreted to suggest that N1303K-CFTR is degraded by a selective autophagic process that requires JB12 and canonical autophagy initiation factors. N1303K-CFTR does not aggregate or damage ER-tubules, and the autophagic intermediates detected are not being consumed by phagophores proposed to engulf damaged ER during ER-phagy (Khaminets, Heinrich et al. 2015, Forrester, De Leonibus et al. 2019). Instead, data are consistent with an ER-membrane containing N1303K-CFTR associating with autophagy initiation machinery present in WIPI1 decorated ER-microdomains. The N1303K-CFTR ER-membrane appears to be modified and converted into ER-connected phagophores that form in association with WIPI1-rings containing docked autolysosomes. These data suggest that ERAD-resistant degradation intermediates of membrane proteins are sorted into ER-connected phagophore membranes. These interpretations are consistent with results from ultra-thin sectioning and electron microscopic studies showing connections between ER tubules and phagophores formed in starved cells (Yla-Anttila, Vihinen et al. 2009, Le Bars, Marion et al. 2014).

### The Conformation of CFTR-Intermediates Controls Triage for ERAD versus ER associated-autophagy

Next, we investigated the role of the conformation of kinetically-trapped intermediates of CFTR in their selection for ER associate-autophagy (Figure 6). The conformation of N1303K-CFTR intermediates was probed with a cysteine (Cys) pair cross-linking assay that monitors the formation of contacts between NBD1 or NBD2 with different CLs identified in high-resolution models of CFTR structure {Serohijos, 2008 #3005}(Figure 6A-C). Cys pairs were introduced at CL/NBD interfaces in Cys-less CFTR with or without N1303K, and cross-linking was conducted to evaluate the folding of CFTR to a conformation that contains the indicated domain interfaces (He, Kota et al. 2013). When a Cys pair was introduced at CL2/NBD2 interface at amino-acid residues C276C/Y1307C, cross-linking of CFTR was detected after treatment of cells with MTS disulfide bond cross-linkers of different spacer arm lengths, M3M and M8M (Figure 6C, lanes 2 and 3, marked with X). VX-809 increased signals that report on the CL2/NBD2 interface within WT-CFTR, accumulating in the low mobility C-form (Figure 6C, lanes 5 and 6). However, in the absence or presence of VX-809, no cross-linked band was detected in N1303K-CFTR that reports on the CL2/NBD2 interface (Figure 6C, lanes 7-12). Similar results were found in N1303D-CFTR (Figure S5B, lanes 1-6), indicating the introduction of a charged residue at N1303 disrupts the NBD2/CL2 interface in a manner that could not be reversed by VX-809.

**Figure 6.**
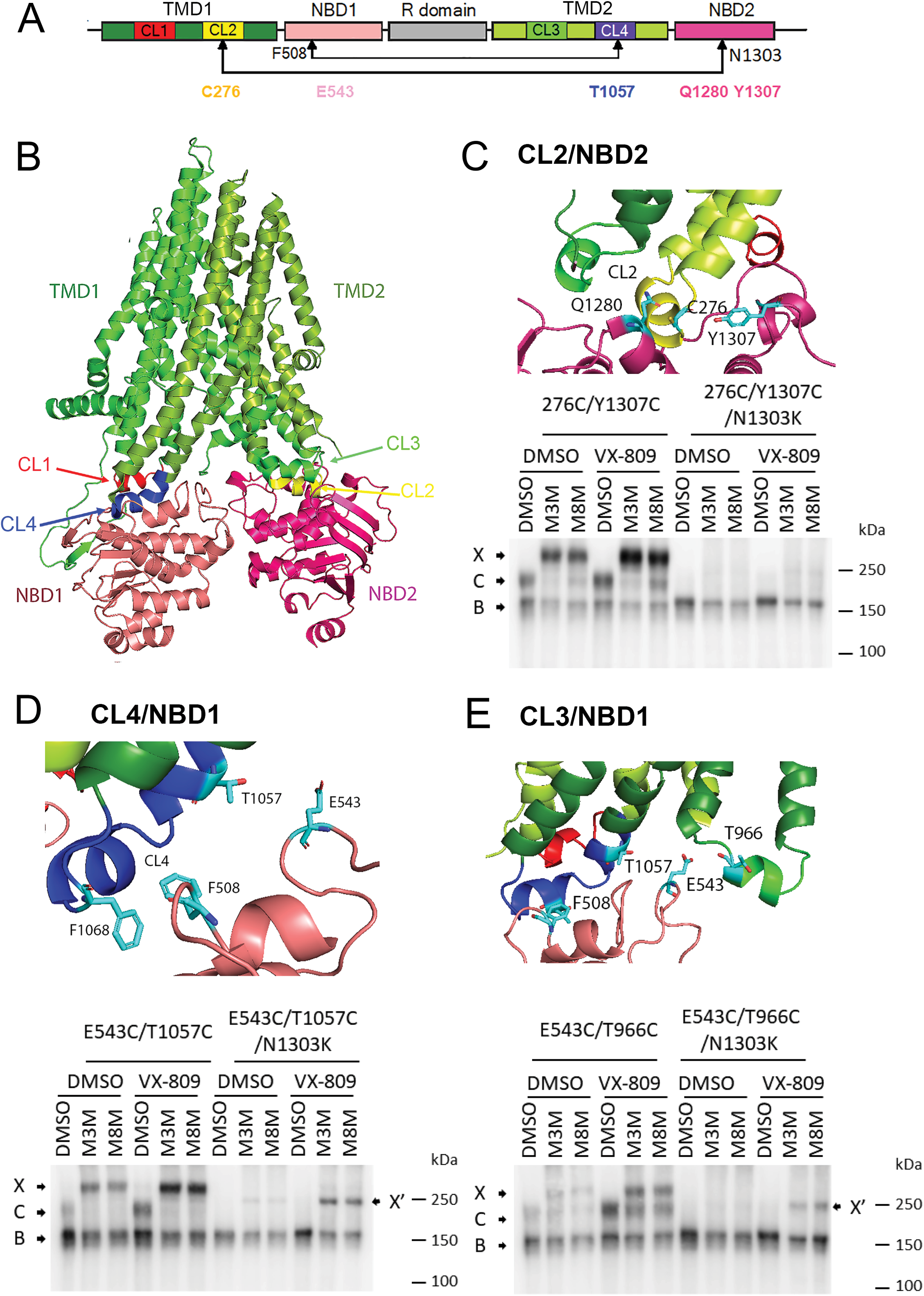
N1303K CFTR folded to a conformation that contains native-like domain:domain contacts between NBD1 and ICL4, but native-like assembly of NBD2 is not detected. (A). Model for domain structure of CFTR containing two nucleotide-binding domains (NBD1 and NBD2), two membrane-spanning domains (MSD1 and MSD2), and a regulatory region (R domain). Each MSD contains two cytoplasmic loops (CL) that form interfaces with the NBDs. F508 in NBD1, N1303 in NBD2, and residues mutated to Cys are marked. (B) The 3-dimensional structure of human CFTR (5UAK) with major domains and CLs labeled. (C-E). Cross-linking of Cys pairs introduced at different CL/NBD interfaces. HEK293 cells were transiently transfected with Cys-less CFTR or Cys-less N1303K CFTR with Cys pairs E543C/T1057C introduced at CL4/NBD1 interface (B), E543C/T966C at CL3/NBD1 interface (C) 276C/Y1307C at CL2/NBD2 interface (D), respectively. At 24 h after transfection, cells were treated with 3 μM VX-809 for 24 h at 27°C. Cells were harvested and resuspended in PBS and incubated with 200 μM M3M, M8M, or an equal amount of DMSO as vehicle control. Cell lysates in SDS-PAGE sample buffer without DTT were subjected to 7.5% SDS-PAGE and Western blot analysis with mAb 596. C, mature complex-glycosylated CFTR; B, immature core-glycosylated CFTR; X, cross-linked mature CFTR; X’, cross-linked immature CFTR. Additional cross-linking experiments are presented in the supplement.

The Cys pair at amino acid residues E543C/T1057C, in NBD1 and CL4, respectively, was acted upon by both MTS cross-linkers M3M and M8M to generate a lower mobility C-form of CFTR (Figure 6D, lanes 2 and 3). The introduction of N1303K into this reporter prevented the C-form of CFTR (Figure 5D, lane 7). Still, a faint cross-linked band was detected slightly above the C-band’s location is seen in extracts of cells treated with M3M and M8M (Figure 6D, lanes 8 and 9, marked with an X’). VX-809 treatment increased the B-form N1303K-CFTR (Figure 6D, lane 10) and the amount of band X’ (Figure 6D, lanes 11 and 12). VX-809 restored the interface between CL3/NBD1 in N1303K-CFTR, as shown with the cross-linking of the E543C/T966C marked with X’ ’Figure 6E).

Sub-domains of N1303K-CFTR assemble into a conformation that has features of folded WT CFTR, but folding steps involving NBD2 are defective and resistant to modulation by VX-809. The combined folding of N1303K-CFTR’s-terminal domains and failure to complete late-stage domain:domain assembly reactions could cause the accumulation of kinetically-trapped intermediates.

### Control of Triage Decisions by the Conformation of N1303K-CFTR

To test the model proposed for conformation-dependent triage of CFTR intermediates, we developed a CFTR reporter that is conditionally degraded in the absence or presence of VX-809 by either ERAD or ERQC-autophagy, respectively (Figure 7A). The disease-causing mutation P67L located in the N-terminus causes global misfolding and degradation of P67L CFTR by ERAD, and VX-809 completely restores the folding and function of P67L CFTR (Ren, Grove et al. 2013). P67L was therefore introduced into N1303K-CFTR to create P67L-N1303K-CFTR. It was then asked: 1. What is the effect of P67L on the triage of N1303K-CFTR (Figure 7B)? 2. Does stabilization of N-terminal domains by VX-809 cause P67L-N1303K CFTR to behave like N1303K-CFTR (Figure 7C). P67L-N1303K-CFTR intermediates exhibited a short half-life and were not detected in autolysosomes. Treatment with VX-809 restored P67L-CFTR folding and accumulation in the C-form and in parallel incubations increases in the stability of P67L-N1303K-CFTR intermediates and their accumulation in autolysosomes was now observed (Figure 7B and 7C). Globally misfolded P67L-N1303K-CFTR goes to ERAD, whereas VX-809 stabilized P67L-N1303K-CFTR is degraded by ERQC-autophagy. At the same time, misfolded CFTR and F508-CFTR are degraded by ERAD in the absence or presence of VX-809. These data demonstrate that its intermediates’ stability dictates the degradative fate of a misfolded and detergent-soluble membrane protein such as CFTR.

**Figure 7.**
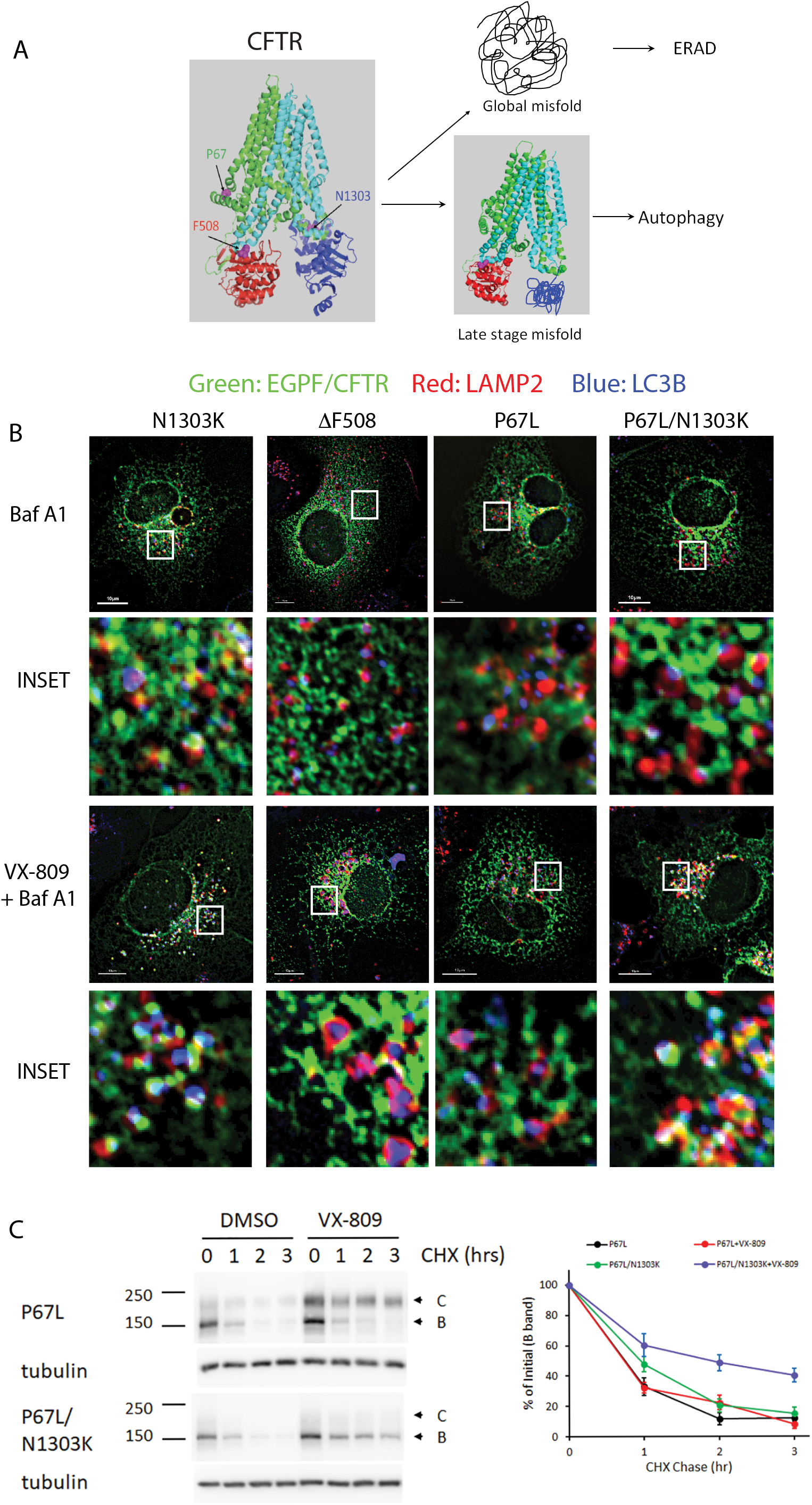
Introduction of N1303K in P67L CFTR renders it to be recognized by autophagy machinery after VX-809 treatment. (A). Model for conformation dependent triage of misfolded CFTR intermediates (B). P67L/N1303L CFTR as a model to test conformation specific triage of CFTR degradation. COS-7 cells were transfected with various EGFP-tagged mutant CFTR. Cells were treated overnight with VX-809 before Bafilomycin A1 treatment for 6 hrs. Immunostaining was carried out with anti-LAMP2 (shown in red) and anti-LC3B (shown in blue) antibodies. (C). CHX chase of P67L and P67L/N1303K CFTR. HEK cells transfected with P67L CFTR or P67L/N1303K CFTR and CHX chase was carried out as described in Figure 2A figure legend.

## Discussion

We report that the mutation N1303K in NBD2 arrests CFTR folding at a late stage and causes kinetically trapped intermediates to accumulate in the ER. N1303K-CFTR intermediates are chaperoned by JB12 and Hsp70 and triaged in a conformation-specific manner for degradation by ERAD or ER associated-autophagy. Data presented suggest that globally misfolded CFTR is degraded by ERAD, whereas biochemically stable intermediates are selected for ER-associated-autophagy. N1303K-CFTR intermediates have long 3-hour half-lives, are soluble in non-ionic detergent, mobile in the ER tubular network, and contain cross-linkable domains interfaces present in folded CFTR. N1303K-CFTR intermediates are stabilized by VX-809, but stabilized N1303K-CFTR does not escape the ER and remains associated with JB12 and additional PQC factors. Disruption of early steps in CFTR folding to cause global misfolding of N1303K-CFTR via the introduction of the P67L mutation reduces the half-life of N1303K-CFTR, and P67L-N1303K-CFTR is not detected in the autophagy system. VX-809 stabilizes P67L-N1303K-CFTR intermediates causing them to be triaged to ERQC-autophagy instead of ERAD. Data reported on N1303K-CFTR support the concept that in energized and biosynthetically active ER membranes, non-toxic intermediates of misfolded membrane proteins are selectively triaged for proteasomal or lysosomal degradation based on their conformation.

CFTR degradation intermediates are now grouped with an expanding collection of secretory and membrane proteins that misfold and accumulate in ERAD-sensitive or ERAD-resistant conformations (Buchberger 2014, Houck, Ren et al. 2014). ERAD requires that misfolded luminal or membrane proteins be unfolded and retrotranslocated to the cytosol, where they are translocated into the narrow opening of proteasomes’ proteolytic chamber. In cases where nascent secretory proteins misfold in the ER-lumen and adopt a retrotranslocation-resistant conformation, the trapped intermediates interact with complexes containing ER-phagy receptors before selective ER-phagy (Fregno and Molinari 2018). We extensively evaluated the role of selective ER-phagy in the conformation-dependent triage of N1303K-CFTR. We did not identify an ER-phagy receptor or soluble autophagy receptor necessary for its conformation-dependent sorting and traffic to autolysosome (Table S1). Since there are a large number of LIR-containing proteins implicated in selective ER-phagy and selective autophagy, it could be that functional redundancy complicates the interpretation of siRNA and CRISPR/Cas9 loss of function experiments. However, we combined gene inactivation with imaging analysis and biochemical studies. We were unable to observe physical or functional interactions between soluble or ER-membrane associated autophagy receptors with N1303K-CFTR that were suggestive of them playing a role in quality control of membrane proteins.

In contrast to results reported with N1303K-CFTR, aggregates of ER-luminal alpha-1-antitrypsin and large assemblies of misfolded collagen are cleared from the ER by a selective ER-phagy process involving FAM134B and the ER exit site protein Sec31 (Forrester, De Leonibus et al. 2019).(Omari, Makareeva et al. 2018). N1303K-CFTR is excluded from ER-exits and does not co-localize with Sec31 or tango, so it is not likely to be degraded by the pathway used for misfolded collagen. The chaperone calnexin binds selective ER-phagy clients in the lumen and functions in a complex with FAM134B to target them for selective ER-phagy. CFTR is a calnexin client, and calnexin facilitates assembly reactions involving C-terminal regions of CFTR, but a role for FAM134B in the degradation of N1303K-CFTR could not be demonstrated (Table S1). These results might be observed because polytopic membrane proteins expose surfaces in the cytosol ubiquitinated when they fail to fold correctly. N1303K-CFTR, like other misfolded forms of CFTR, is ubiquitinated, which could permit cytosolic ubiquitin chain binding autophagy receptors to usurp requirements for ER-phagy receptors. However, we could not demonstrate a role for p62, NBR1, FKBP8, or Tollip in the conformation-dependent sorting of misfolded membrane proteins to the autophagy initiation sites when ER-tubules contain N1303K-CFTR associate with WIPI1-rings (Table S1). Mechanisms for the conformation-dependent selection of misfolded and detergent soluble forms of CFTR for selective ER-associated autophagy, therefore, appear different than those demonstrated for ERAD-resistant luminal aggregates and aggregates of cytosolic proteins.

Triage of nascent CFTR between folding and degradation is mediated by JB12, Hsp70, and ERQC-E3 ligases such as RMA1/RNF5 and CHIP (Meacham, Lu et al. 1999, Meacham, Patterson et al. 2001, Younger, Chen et al. 2006, Grove, Fan et al. 2011). N1303K-CFTR is present in complexes with JB12 and Hsp70, and siRNA depletion of JB12 hinders degradation of N130K-CFTR and blocks the accumulation of N1303K-CFTR in autolysosomes. The overexpression of dominant-negative QPD JB12 also prevents N1303K-CFTR from being degraded by ERAD or autophagy. QPD-JB12 can bind clients, but cannot interact with Hsp70, so QPD-JB12 can bind but fails to efficiently release clients (Grove, Fan et al. 2011). JB12 is not required for basal autophagy, so a functional interaction of JB12 and Hsp70 appears needed to deliver N1303K-CFTR to the ER’s autophagy machinery. JB12 is detected in complexes with Beclin-1, and complex formation between JB12 and Beclin-1 is stimulated by ERQC-autophagy clients (Houck, Ren et al. 2014). Therefore, stable client binding to JB12 could trigger the focal nucleation of autophagy initiation machinery (Joo, Dorsey et al. 2011, Pickles, Vigie et al. 2018) and facilitate the observed intramembrane sorting of its ERAD-resistant N1303K-CFTR to omegasomes. However, additional studies are required to determine the mechanism by which JB12 and Hsp70 assist in the sorting of stable degradation intermediates of membrane proteins to the autophagy initiation sites within the ER.

To define the pathway for N1303K-CFTR intermediates from the ER to the autolysosome, we carried out a combination of fixed and live-cell imaging at normal and super-resolution. In particular, we sought to determine if N1303K-CFTR entered autolysosomes via an ER-phagy like mechanism in which ER-subdomains containing it are consumed by phagophores or a focal mechanism involving direct entry of N1303K-CFTR into membranes for ER-derived autolysosomes (Houck, Ren et al. 2014). To test the latter possibility, the association of ER-microdomains containing N1303K-CFTR or F508del-CFTR with autophagy initiation site machinery (WIPI1, LC3B, FIP200) that localize within omegasomes and phagophores was evaluated. ER membranes enriched in N1303K-CFTR and F508del-CFTR were detected on the perimeter of WIPI1 decorated ER membranes, but only N1303K-CFTR was detected in ER-associated membranes which were decorated with the phagophore marker LC3B. Likewise, N1303K-CFTR was also seen in WIPI1 enriched foci with the ULK1 subunit FIP200. In contrast, JB12 was excluded from ER-tubules that are decorated with LC3b and/or associate with WIPI1 and FIP200. It was common for LAMP1 enriched lysosomes to be detected in association with ER membranes containing N1303K-CFTR that extend from the ER surface. There were also N1303K-CFTR and LC3B containing foci in the 50 nm size-range that were sandwiched between the ER and docked lysosomes. Such foci appear to represent autophagosomes in the act of docking with lysosomes. These imaging data are consistent with the concept of ER associated-autophagy involving the focal activation of selective autophagy in response to the accumulation of ERAD-resistant intermediates of N1303K-CFTR.

N1303K-CFTR is the second most common CF causing allele, and it was hoped that folding modulators developed to restore the F508del-CFTR function could also fix the N1303K-CFTR function. The Class I modulator VX-809 does increase the accumulation of N1303K-CFTR, but not proper folding. Treatment with VX-809 alone, or in combination with the tool compounds in the Class II modulator family, or VX-441, which is used in combination with Class I modulators clinically, does not restore N1303K-CFTR function in heterozygous or homozygous native human airways cells from CF patients (Veit, Roldan et al. 2020). However, we were able to fix the folding of a small fraction of N1303K-CFTR in cells that are cultured at low temperatures. Reconstituted N1303K-CFTR channels isolated from the plasma membranes of cells cultured at low-temperature and assayed over a range of temperatures had the same conductance as WT CFTR. Rescued N1303K-CFTR channels were, however, thermodynamically unstable with a dramatically lower open probability than WT CFTR. Notably, the CFTR channel potentiator VX-770 was found to enhance the open probability of N1303K-CFTR at temperatures between 20 and 37 °C, but rescued channels remained unstable.

Folding/assembly of NBD2 is a slow step in CFTR biogenesis that involves cooperative conformational changes that result from dimerization of NBD2 with NBD1 and reorientation of transmembrane spans and CLs (Wang, Wu et al. 2010, Kirk and Wang 2011, Veit, Xu et al. 2018). Defects in the folding of N1303K-NBD2 could block the folding progression of partially assembled intermediates for several reasons. Class I and Class II modulators were developed to overcome defects in F508del-NBD1 folding/assembly, and they are also able to suppress functional defects in a subset of non-F508del CFTR mutants. Yet, rare CFTR alleles that respond to Trikaftor have common biogenic defects similar to those caused by F508del (Middleton, Mall et al. 2019). Such defects are suppressed allosterically by stabilizing N-terminal regions of CFTR and promoting intramolecular domain: domain assembly reactions involving F508del-NBD1 (Ren, Grove et al. 2013, Veit, Avramescu et al. 2016). Clinically relevant restoration of the N1303K CFTR function might be achievable through the development of modulators that rescue defects in NBD2 misfolding and/or assembly.

## Materials and Methods

### Reagents and antibodies

Reagents and antibodies were purchased from the following suppliers: HRP-conjugated goat anti-mouse IgG (31430) and goat anti-rabbit IgG (31461), Alex Fluor 488 goat anti-mouse IgG (A11029), Texas Red goat anti-mouse IgG (T-862), and goat anti-rabbit IgG (T-2767), Alexa Fluor Plus 647 goat anti-mouse IgG (A32728) and goat anti-rabbit IgG (A32733) from Thermo Fisher Scientific. Anti-beclin-1 (NB500-249), anti-FAM134B (NBP2-55248) and anti-Sec62 (NBP1-84045) from Novus Biologicals. anti-calnexin (C4731), anti-HA (H9658), anti-LC3B (L7543), anti-Myc (C3956), and anti-tubulin (T-9026) from Sigma-Aldrich. Anti-ERGIC53 (ALX-804-602) from Enzo Life Sciences. Anti-GABARAP+L1+L2 (ab109364) and anti-GM130 (ab52649) from Abcam. Anti-LAMP2 (555803) and anti-Sec31A (612351) from BD Pharmingen. Anti-p62 (H00008878-M0162) from Abnova. Anti-VMP1 (12978s) from Cell Signaling. DC Protein Assay (500-0119) and Clarity Western ECL Substrate (170-5061) from BioRad. Effectene Transfection Reagent (301427) from Qiagen. Lipofectamine RNAiMAX Reagent (100014472) from Thermo Fisher Scientific. Methanethiosulfonate (MTS) cross-linking reagents M3M (1,3-propanediyl bismethanethiosulfonate) and M8M (1,5-pentanediyl bismethanethiosulfonate) from Toronto Research Chemicals. ProLong Diamond antifade mountant without (P36970) or with DAPI (P36971) from Thermo Fisher Scientific.

### Cell culture and transfection

African green monkey kidney cells (COS-7 ATCC CRL-1651), human embryonic kidney cells (HEK293, ATCC CRL-1573), and baby hamster kidney cells (BHK-21, ATCC CCL-10) were cultured in DuDulbecco Eagle minimum enriched medium (DMEM) supplemented with 10% fetal bovine serum (Sigma-Aldrich, St. Louis, MO) and 10U/ml penicillin, 10μg/ml streptomycin (GIBCO, Carlsbad, CA) at 37°C, 5% CO_2_. Human lung tissue was procured under The University of North Carolina Office of Research Ethics Biomedical Institutional Review Board Approved Protocol No. 03–1396. Primary HBE cells were harvested and cultured using established procedures previously described in detail (Ref).

Plasmid transfections in cell lines were performed using Effectene Transfection Reagent following the product instruction. For knockdown experiments, all siRNAs were purchased from Thermo Fischer Scientific. For some target genes, two siRNAs to the same target gene were used to increase knockdown efficiency. COS-7 or HEK293 cells that were seeded into a 6-well plate at 3×10^5^ cells/well for 20 hr. were transfected at five pmol per well of siRNA with Lipofectamine RNAiMAX. Twenty hours post-transfection, the cells were split again to the appropriate density for transfection of plasmids using Effectene Transfection Reagent. siControl siVMP1 and sip62 sequences:

### Protein sample preparation for SDS-PAGE and Western blotting

COS-7 cells or HEK293 cells were harvested from plates with citric saline (135mM KCl, 15 mM Sodium Citrate), and cell pellets were resuspended in PBST lysis solution (1% (v/v) Triton X-100 in phosphate-buffered saline) supplemented with one mM PMSF, 1x complete protease inhibitor cocktail (Roche, Basel, Switzerland). 2x Laemmli sample buffer without reducing agent was added, and the total lysates were homogenized by sonication. Protein concentrations were measured using detergent compatible DC Protein Assay Reagent, and 50 mM DTT was added before samples were resolved by SDS-PAGE and proceeded to Western blot analysis. Protein signal was detected by Clarity™ western ECL substrate and visualized by LAS680 imager (GE Life Sciences, Pittsburgh, USA). Images were analyzed by ImageJ (NIH, USA) and quantified with ImageQuant (GE Life Sciences, Pittsburgh, USA).

### Co-immunoprecipitation assay

HEK293 or COS-7 cell lysates in PBST solution (1% Triton X-100 in PBS) were centrifuged at 100,000 g, at 4 °C, for 10 min to remove any insoluble fraction. Protein concentrations were measured using DC Protein Assay, and samples of an equal amount of proteins were used for immunoprecipitation analysis. To pull-down proteins interacting with CFTR, mAb596 antibodies cross-linked to Protein G agarose beads (Invitrogen) were incubated with cleared lysate for 1 hr. at 4 °C. Beads were washed 3x with PBST lysis buffer. Bound proteins were eluted from beads via incubation with 2x SDS-PAGE sample buffer without reducing agent at 37°C for 15 min. Anti-FLAG M2 beads (Sigma-Aldrich, St. Louis, USA) were used to pull-down proteins interacting with FLAG-tagged proteins. After washing, proteins bound were eluted with FLAG peptide. Eluted proteins were subjected to SDS-PAGE and Western blot analysis as described above.

### Cycloheximide chase

To determine the steady-state level of different CFTR mutants, HEK293 cells transfected and grown overnight prior to incubation with 5μg/ml cycloheximide (CHX; Sigma-Aldrich, St. Louis, MO) in the presence and absence of 15μM CQ (Sigma-Aldrich, St. Louis, MO) or 10μM bortezomib (LC Laboratories, Woburn, MA). Cells were harvested at indicated time points, and protein samples were prepared for Western blot analysis as described above.

### Triton X-100 solubility assay

Harvested cell pellets were lysed in PBST (1% Triton X-100 in PBS) and separated into soluble and insoluble fractions by centrifugation at 20,000xg for 30 min at 4°C. The insoluble fraction was resuspended in a 2x sample buffer with an equal volume of soluble fraction and sonicated. Total lysates, soluble fractions, and insoluble fractions were resolved with SDS-PAGE and subjected to Western blot analysis.

### Disulfide cross-linking in whole cells

Cross-linking of Cys pairs introduced in Cys-less CFTR expressed in HEK293 cells was performed essentially as previously described (Serohijos, Hegedus et al. 2008). Briefly, HEK cells transiently transfected with Cys-less CFTR constructs with Cys pairs introduced at CL/NBD interfaces were grown in 6 well plates. The cell was harvested in citric saline, the cell pellet was washed in PBS and then resuspended in 30 μl PBS. 10 μl of cell suspension was mixed with 20 μl PBS with DMSO as vehicle control or PBS containing 300 μM MTS cross-linker to yield a final concentration of 200 μM of bifunctional cross-linkers M3M (1,3-propanediyl bismethanethiosulfonate), or M8M (1,5-Pentanediyl bismethane-thiosulfonate) (Toronto Research Chemicals). After 15 min incubation at room temperature, the cross-linking reaction was stopped with 120 μl 1.5x Laemmli sample buffer without DTT. After sonication, 30 μl of the samples were loaded on 7.5% SDS-PAGE, and anti-CFTR mAb 596 was used for Western blotting.

### Immunofluorescence microscopy

COS-7 cells were used for most of the microscopy study due to their flat and spread out nature, making them ideal for resolving intracellular organelles. COS-7 cells were grown on thin coverslips (Thermo Fisher Sci. 12541A) in 6 well plates. After transfection and treatment, cells were fixed at −20 °C for 5 min with pre-cooled methanol. Cells were rehydrated by washing 3x in PBS and blocked with 3% BSA in PBS for 30 min, before immunostaining with various antibodies. Goat anti-mouse or goat anti-rabbit secondary antibodies with different fluorescent tags were used to detect proteins of interest. Coverslips were mounted onto slides using ProLong Diamond antifade mountant without (P36970) or with DAPI (P36971) (Invitrogen). Slides were allowed to dry in the dark at room temperature for at least 12 hours before imaging with an IX81 motorized inverted microscope (Olympus, Center Valley, PA) equipped with standard DAPI (blue), FITC (green), TRITC (red), Cyan (far red) filter cubes. STED images were collected with a Leica TCS SP8 3X equipped and a FOV scanner. Acquired images were deconvoluted using CellSens Software (San Jose, CA, USA) to remove background noise.

### Live cell imaging

COS-7 cells were cultured in 35mm glass-bottom dishes (MatTek P35G-1.5-10-C). After overnight transfection with fluorescent-tagged proteins, cells were rinsed with FluorBrite DMEM (Gibco Life Sci.) and kept in the same medium during drug treatment and imaging. Live cell imaging was carried out using an Olympus FV3000RS Confocal Microscope equipped with a humidified incubation chamber set to 37 °C. Images were acquired and deconvoluted using CellSens software (Olympus).

### Measurement of CFTR activity in USSING chambers

Ion transport measurements with HBEs were performed in modified Ussing chambers (Physiologic Instruments) under voltage clamp-conditions using Acquire & Analyze (version 2.3) software (Physiologic Instruments). Reference measurements [potential difference (*P*_d_)] and transepithelial resistance (*R*_t_) were obtained for each culture. Short circuit current (*I*_sc_) was measured every 20 s and recorded digitally after bilateral equilibration at pH 7.4 for 10 min in 5 ml of pH 7.4 Krebs bicarbonate Ringers (KBR; 115 NaCl mM, 25 NaHCO_3_ mM, 2.4 K_2_HPO_4_ mM, 1.2 CaCl_2_ mM, 1.2 MgCl_2_ mM, 0.4 KH_2_PO_4_ mM, and five D-glucose mM). Chamber temperature was maintained at 36°C ± 1°C by a circulating water bath, and KBR was bubbled with 95% O_2_-5% CO_2_ throughout the experiment.

Agonists and inhibitors used to modulate CFTR activity were purchased from Sigma Chemical. In the presence of amiloride (100 μM, apical), forskolin (10 μM) was applied bilaterally to induce cAMP activation of CFTR and VX-770 (5 μM, apical) was applied to further activate CFTR. Currents detected were sensitive to the CFTR channel blocker CFTRinh-172 (10 μM apical). As an internal control, UTP (100 μM) was used to activate calcium-activated chloride channels (CaCC) at the conclusion of the experiment. Data were exported and analyzed in Microsoft Excel, and slope values were subtracted for accuracy when necessary. Mean, SD, and SE measurements were calculated for each culture. Statistical analysis was performed by an unpaired two-tailed Student’ s-test. *P* < 0.05 was considered to indicate statistical significance. Line and Bar graphs were created using Origin 8.6 software (OriginLab).

### Membrane isolation and planar bilayer based single-channel recording

BHK cells stabling expressing N1303K CFTR with or without stabilizing mutations H1402S and ΔRI/2PT/H1402S were grown to 90% confluency, and cells were moved to a 27 °C CO_2_ incubator and grown with three μM VX-809 for 24 hrs. Cells were harvested and homogenized on ice in 10 mM HEPES, pH7.2, one mM EDTA containing a protease inhibitor cocktail. Centrifugation at 3000 g for 5 min removed nuclei and undisrupted cells. The supernatant was centrifuged at 100,000 g for 30 min to pellet membranes, which were then resuspended in phosphorylation buffer (10 mM Hepes, pH 7.2 containing 0.5 mM EGTA, two mM MgCl_2_, and 250 mM sucrose).

Planar lipid bilayers were prepared by painting a 0.2 mm hole drilled in a Teflon cup with a phospholipid solution in n-decane containing a 3:1 mixture of 1-palmitoyl-2-oleoyl-sn-glycerol-3-phosphoethanolamine and 1-palmitoyl-2-oleoylsn-glycerol-3-phosphoserine (Avanti Polar Lipids). The lipid bilayer separated 1.0 ml of solution (*cis* side) from 5.0 ml of solution (*trans* side). Both chambers were magnetically stirred and thermally insulated. Heating and temperature were controlled with a Temperature Control System TC2BIP (Cell Micro Controls). CFTR ion channels were transferred into the preformed lipid bilayer by spontaneous fusion of membrane vesicles containing CFTR. To maintain uniform orientation and functional activity of CFTR channels, two mM ATP, 50 nM PKA, and membrane vesicles were added to the *cis* compartment only. All measurements were done in symmetrical salt solution (300 mM Tris/HCl; pH 7.2; 3 mM MgCl_2_ and 1 mM EGTA) under voltage-clamp conditions, using an Axopatch 200B amplifier (Axon Instrument/Molecular Device). The membrane voltage potential of −75 mV is the difference between *cis* and *trans* (ground) compartments. The output signal was filtered with an 8-pole Bessel low-pass filter LPBF-48DG (NPI Electronic, Tamm, Germany) with a cut-off frequency of 50 Hz, digitized with a sampling rate of 500 Hz, and recorded with pClamp9.2software. Origin 75 software (Origin Lab Corp., Northampton, MA, USA) was used to fit all-points histograms (pClamp 9.2, Axon Instruments) by multi-peak Gaussians. Single-channel current was defined as the distance between peaks on the fitting curve and used for the calculation of the single-channel conductance. The probability of the single-channel being open (*P*_o_) was calculated as a ratio of the area under the peak for the open state to the total area under both peaks on the fitting curve.

## Acknowledgments

This work was supported by grants from the Cystic Fibrosis Foundation (CYR18XX0, BOUCHE19R0, GENTZS19I0) and NIH (GM056981 and P03DK065988). FRAP and STED Images were collected with the assistance of the UNC Hooker and Neuroscience Imaging Core facilities. Susan Boyle assisted with the culture of primary human bronchial epithelial cells.

