## Supplementary material for "DNAJB12 and Hsp70 Facilitate the Conformation Specific Degradation of Arrested N1303K-CFTR Intermediates by ER Associated-Autophagy": Sup Fig 1-4, TS1

### Figure S1

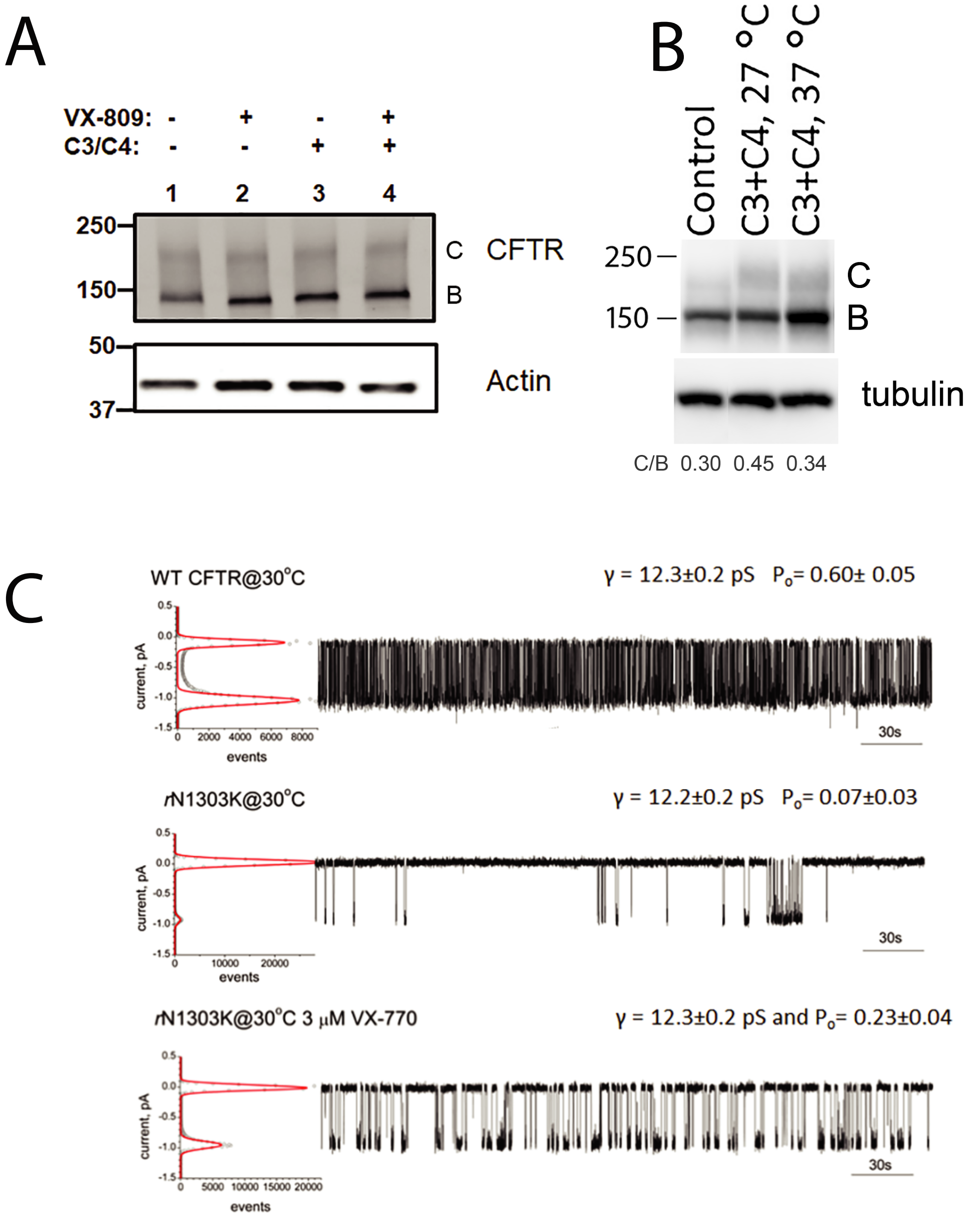


Figure S1 associated with Figure 1. Analysis of N1303K-CFTR expression and function (A) The effect of small molecule folding modulators on steady-state levels of full length CFTR polarized native N1303K/W1282X human bronchial epithelia (HBE). Signals for truncated W1282X CFTR are not detected and not shown. B. Steady-state levels of N1303K CFTR stably expressed in BHK cells use for enrichment of CFTR channels in membrane fractions of cells. Incubations with modulators and at different temperatures were as described in the legend to Figure 1 and in the materials and methods. (C) Single channel function of the WT and rescued N1303K (rN1303K) CFTR in the lipid bilayer at 30 0C. 5 independent experiments of total 42 minutes, 4 independent experiments of total 28 minutes duration, and 3 independent experiments with total 12 minutes duration were used to calculate γ and Po of WT CFTR, rN1303K and rN1303K+VX-770, respectively, at 37 °C.

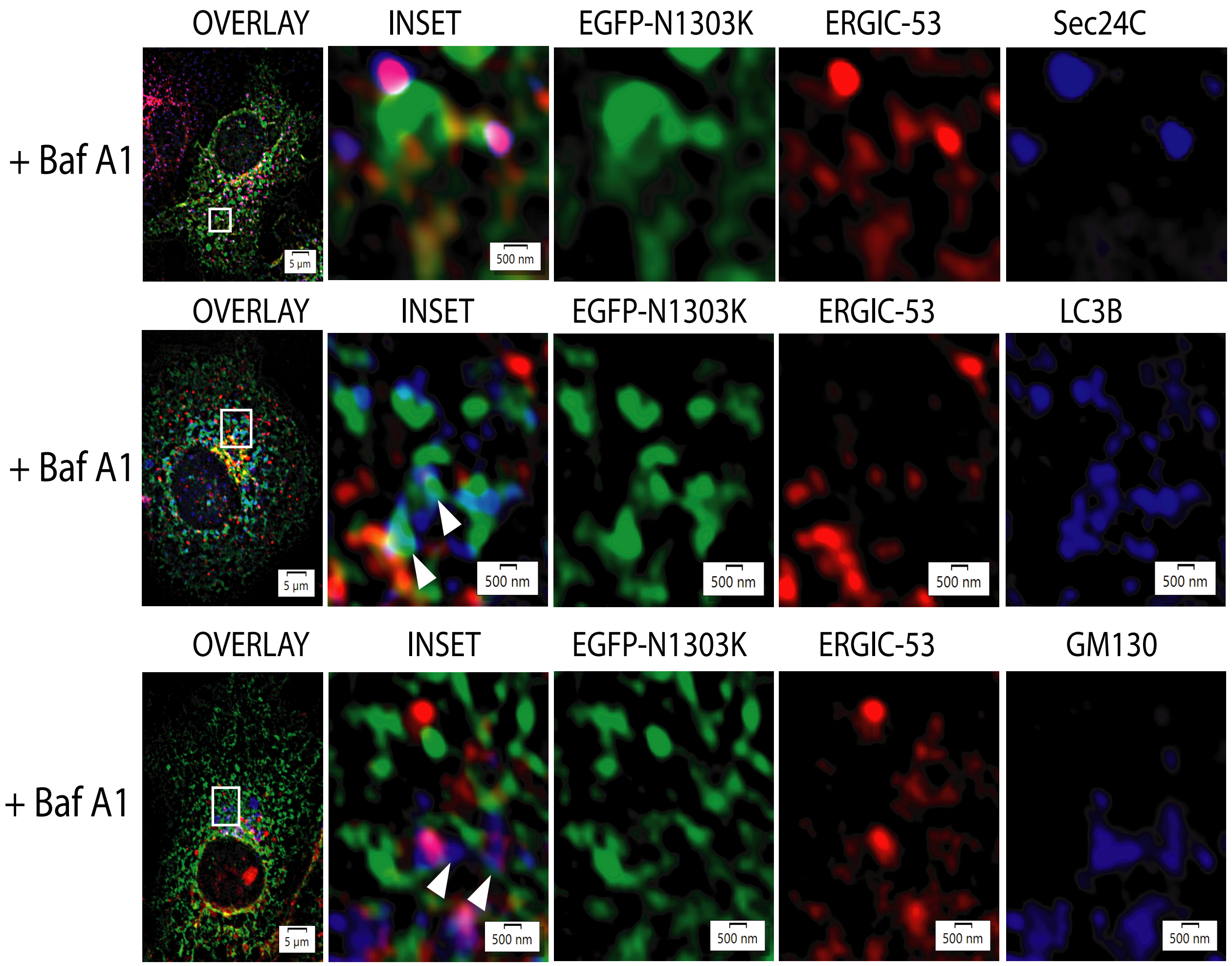


Figure S2 associated with Figure 2. N1303K-CFTR localization in cells with markers for ER exit sites (Sec24), the ER intermediate compartment (ERGIC53), golgi apparatus (GM130), and phagophores (LC3B). COS-7 cells were transiently transfected with N1303K CFTR. 18 hrs. after-transection and 6 hrs. prior to fixation cells were treated with bafilomycin for 6-hours. Immunostaining was carried out on methanol fixed cells with anti-GFP, and with antibodies to endogenous ERGIC-53, GM130, LC3B, and Sec24. Data was collected and process as described for Figure 2 panel B.

Figure S3

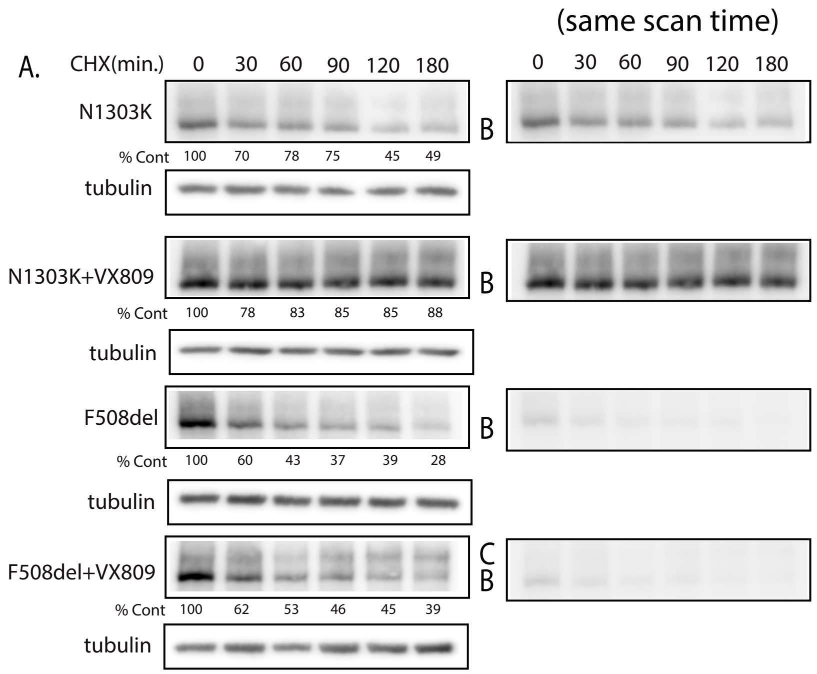


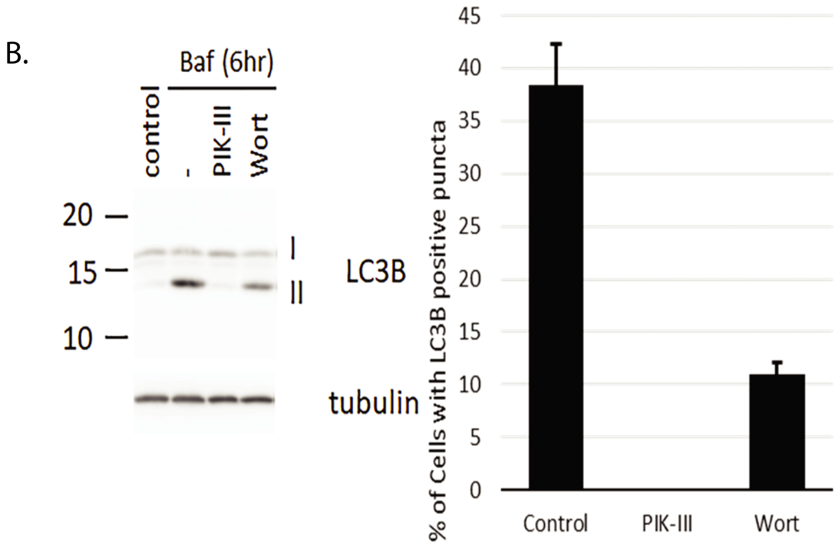


Figure S3 associated with Figure3. A. Steady-levels of N1303K-CFTR in normalize (left panel) and raw (right panels) in cells grown under conditions where overexpressed JB12 and RMA1.
B. Impact of PIK-III on processing of LC3BI to LC3B2 in cell where it blocked entry of N130K-CFTR into autolysosomes is blocked. Indicated forms of CFTR were expressed in the absence or presence of VX-809 that was added to cell cultures 6-hours prior to harvest and analysis by western blot.
Cells were grown and treated as described in the legend to figure 3 and the materials and methods section.

**Figure S4**

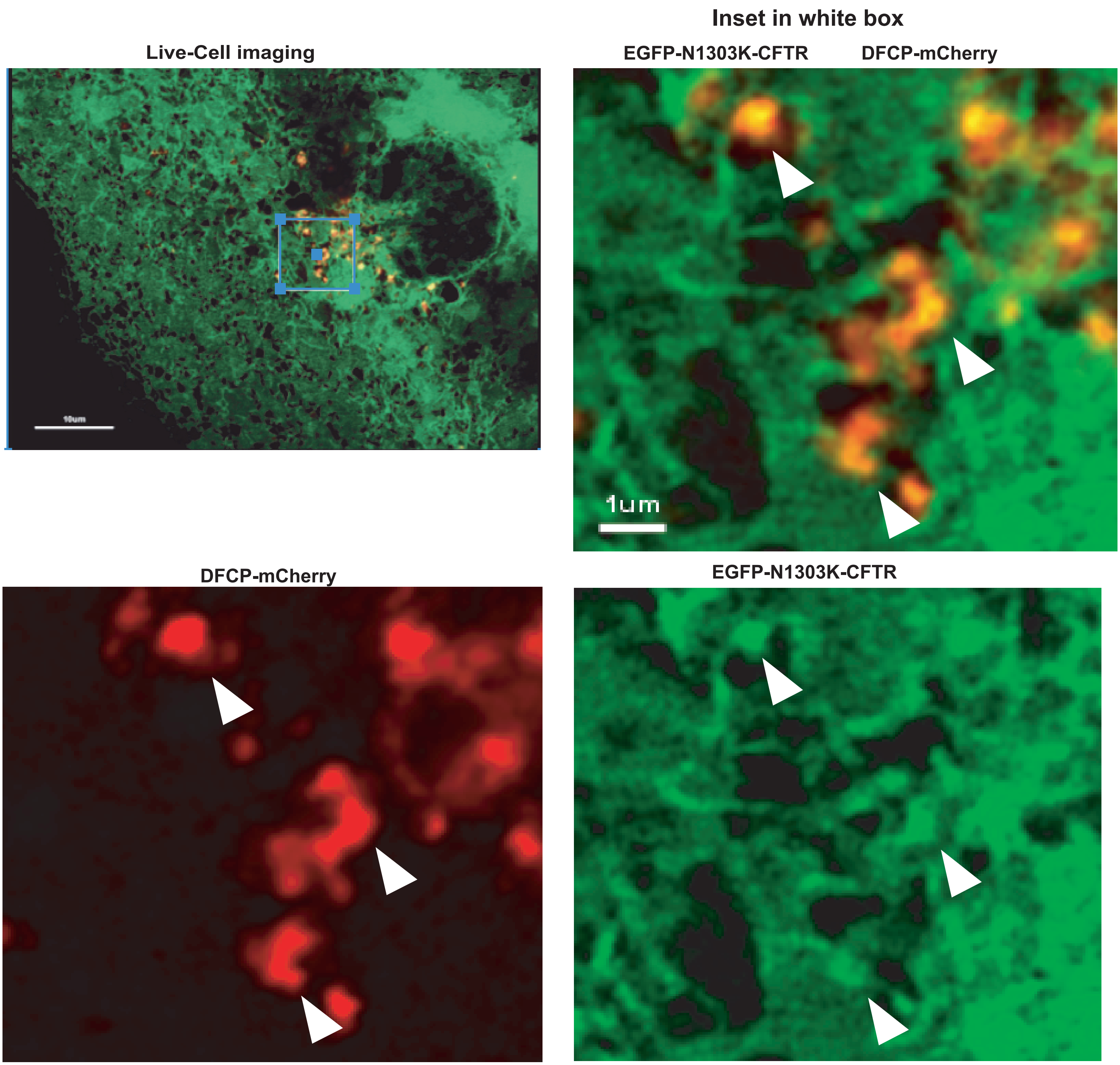


Figure S4 associated with Figure 5. Localization of EGFP-N1303K-CFTR with DFCP-mCherry in live COS-7 cells. Images were taken of live COS-7 cells that did not receive chemical treatment with an Olympus 880 with Airyscan microscope. The inset illustrates the association of ER-tubules containing N1303K-CFTR with those decorated by DFCP. Arrows denote examples signals from EGFP-N1303K-CFTR are detected in ER-tubules that associate with the surface of DFCP-mCherry structures and appear to expand and extend out into the cytosol.

Table S1. Impacts of Inactivation of Autophagy Machinery or ER-homeostasis Factors on Triage of N1303K-CFTR to ERQC-Autophagy.

| Component | Cellular Function/Action | Manipulation | Effect on N1303K-CFTR localization with LAMP1/LC3B | Comments |
|---|---|---|---|---|
| **FAM134B** | ER-Phagy | siRNA/CRISPER | None | Not detected in COS-7 |
| **TEX264** | ER-Phagy | siRNA | None | Not present in autolysosomes with N1303K-CFTR |
| **CCPG1** | ER-Phagy | siRNA | None | Not present in autolysosomes with N1303K-CFTR |
| **RTN3** | ER-Phagy | siRNA | None | Not present in autolysosomes with N1303K-CFTR |
| **RTN4** | ER-Phagy | siRNA | None | Not present in autolysosomes with N1303K-CFTR |
| **ATL3** | ER-Phagy | siRNA | None | Not present in autolysosomes with N1303K-CFTR |
| **Beclin-1** | Autophagy Initiation | siRNA<br>DN-Mutant | Blocks | Localized in omegasomes with N1303K-CFTR |
| **ULK1** | Autophagy Initiation | siRNA<br>DN-mutantInhibition by MRT68921 | Blocks | Localized in omegasomes with N1303K-CFTR |
| **VSPS34** | Autophagy Initiation | Inhibition by PIK3-III | Blocks | |
| **VMP1** | Autophagy Initiation | siRNA/CRISPER | Difficult to Evaluate | Causes ER-stress due to defects in Ca2+ metabolism |
| **SEC62** | Translocon Associated | siRNA | Difficult to Evaluate | SiRNA causes general defects in membrane protein biogenesis. Not present in omegasomes or autolysosomes with N1303K-CFTR |
| **DNAJB12** | Transmembrane Hsp40 | siRNA | Blocks | Detected in complexes with misfolded CFTR Prolonged association with N1303K-CFTR |
| **DNAJB14** | Transmembrane Hsp40 | siRNA | None | Relative of DNAJB12, expressed at low levels in ER |
| **P62** | Autophagy Receptor | siRNA | None | Detected in autolysosome with N1303K-CFTR. Not detected in ER with N1303k-CFTR |
| **TOLLIP** | Autophagy Receptor | siRNA | None | Detected in autolysosome with N1303K-CFTR |
| **FKBP8** | Hsp70 co-factor and Autophagy-receptor | siRNA | None | In immuneprecipitates with JB12, Hsp70 and N1303K-CFTR. Not present in omegasomes with N1303K-CFTR. |
| **NBR1** | Autophagy Receptor | siRNA | None | |

Experiments were conducted in HEK293, COS-7 and HELA cells and similar results were obtained in each cell line. FAM134B is not expressed in COS-7 cells but is detected in HELA and HEK293 cells. The end point for assays is detection of N1303K-CFTR in LAMP1/LC3B positive foci.
